# Micro-C in *Solanum* Uncovers Conserved Genome Folding and Epigenetically Defined Loops with Bifunctional Enhancer-Silencer Activity

**DOI:** 10.1101/2025.10.16.682740

**Authors:** Shdema Filler-Hayut, Anders Sejr Hansen

## Abstract

Long-range chromatin loops are key to genome organization, yet their landscape in plants remains poorly defined. Here, we generated a 1.45-billion-contact Micro-C map of cultivated tomato (*Solanum lycopersicum*) that resolves ∼4,600 loops, seven times more than previous Hi-C data. Loop anchors segregate into two epigenetic classes: 1) promoter-centered loops with anchors enriched for RNA polymerase II, accessible chromatin, active histone modifications, and 2) loops with anchors enriched for Polycomb-linked or heterochromatic signatures. Both classes strongly repel young Gypsy retrotransposons, while retaining older copies directly at the anchor, revealing an age-selective filter. Unlike mammalian genome organization, loop anchors rarely overlap insulation boundaries, indicating that loop and domain formation are largely independent. Cross-species Micro-C in two wild tomatoes shows conserved anchor positions despite rapid sequence turnover. Integrating cross-species transcriptomics further reveals a dual regulatory role for promoter-anchored loops: they can enhance expression when paired with activating chromatin or poise/repress it via H3K27me3-rich, distal anchors. These data provide a high-resolution framework for interpreting plant gene regulation in 3D and a resource to probe how looping shapes trait variation and can be leveraged in breeding.

## Introduction

Eukaryotic genomes are folded in a series of nested layers^1^. The DNA double helix first wraps around histones to form nucleosomes, which are then packed into chromatin fibers. These fibers curve into chromatin loops that juxtapose distant genomic elements. Loops aggregate into self-interacting domains. Separately, genomic regions with similar epigenetic states segregate into euchromatic *A* and heterochromatic *B* compartments. At the largest scale, each chromosome occupies its own territory. In vertebrates, domains called Topologically Associating Domains (TADs)^2^, form through dynamic process where cohesin-mediated loop extrusion is halted by pairs of inward-facing CTCF sites (for review^3^). These borders do far more than delineate structure: they are insulating genomic regions by limiting enhancer reach (for review^4^), synchronizing gene activity^5,6^, shaping replication timing^7^, and steering DNA repair^8^. When insulation fails, developmental disorders and cancers can arise^9,10^.

Plants retain the full cohesin complex^11^ but lack a CTCF homologue. Yet, Hi-C maps reveal cohesin-dependent^12^, CTCF-independent interaction domains in Arabidopsis^13^, maize^14^, rice^15^, tomato^16^ and other plant species. These domains have also been shown to insulate genes like canonical vertebrate TADs (for review^17,18^). Like their animal counterparts^19^, plant domains are hierarchically nested, with finer levels emerging as map resolution improves^20^. Because the vertebrate TAD definition hinges on convergent CTCF sites^21^, which are absent in plants, we avoid the term “TAD” here. Instead, we refer to the insulated boundaries of domains as Insulation Valleys (IVs).

By contrast, high-resolution, genome-wide characterization of chromatin loops in crop plants, and their regulatory roles, has lagged behind. In mammals, such loops are known to juxtapose enhancers and promoters to fine-tune transcription (for review^22^). In plants, however, enhancer-like bidirectional unstable RNAs, the hallmark of many mammalian enhancers are uncommon; when present they often arise from promoters, and STARR-seq indicates that regions initiating stable transcripts show stronger enhancer activity than sites with unstable or bidirectional RNAs^23^. Resolving loop classes, their chromatin environments, their relationship to IVs, and their connection to transcription therefore requires loop-scale assay that exceed standard Hi-C resolution^24,25^.

Across plants, reports link chromatin loops to enhancing or repressive expression patterns, but these outcomes largely track the method used. *Arabidopsis thaliana* Micro-C resolves short promoter–terminator “gene loops” enriched at highly expressed genes and near super-enhancers shows enrichment of MED12, MORC, and SWI/SNF (SAS) rather than RNAPII or H3K27me3^26^, consistent with activation-leaning contexts. By contrast, an Arabidopsis Hi-C study identified hundreds of long-range H3K27me3/PRC2-dependent loops that frequently bridge tandem or metabolic gene clusters and are conserved across species, aligning with repressive or poised states^27^. An open-chromatin–anchored TAC-C approach in cereals recovered loops in activating environments and related loop presence to homoeolog expression bias in wheat^28^, while cotton Hi-C across 12 tissues cataloged mostly tissue-specific loops with active marks at anchors; short-range, gene-linked loops were more conserved and A-compartment-biased, whereas very long or highly networked loops associated with fewer active marks and lower expression^29^. These method-dependent trends, open-chromatin-focused assays favoring activating contacts, and conventional Hi-C often capturing Polycomb-linked long-range contacts-underscore that a high-resolution, structure-first, chromatin-state agnostic approach, such as deep Micro-C is needed to reconcile these results and define when plant loops associate with activation versus repression.

Building a high-resolution loop catalog in tomato would (i) resolve loop classes and their chromatin states, distinguishing promoter-centered from Polycomb/heterochromatin-linked architectures; (ii) map and prioritize distal cis-regulatory elements genome-wide, providing candidates for enhancers and repressors; and (iii) quantify conservation and divergence of looped interactions across Solanum, illuminating natural regulatory circuitry with direct relevance for trait variation and accelerated breeding.

Here we present deep Micro-C maps for cultivated tomato that deliver a nucleosome-scale view of a model fruit-crop genome. We use IVs primarily as a benchmark, confirming prior plant observations and method performance, while focusing on loops: their epigenetic signatures, organization relative to other 3D features, evolutionary conservation, and transcriptional function. Two epigenetically distinct loop-anchor classes emerge (promoter-centered vs. Polycomb/heterochromatin-linked), and both anchor types repel young Gypsy retrotransposons while tolerating older copies, consistent with age-selective TE filtering. Unlike mammals, loops seldom coincide with boundaries, suggesting boundary-independent loop formation. Comparative Micro-C with two wild tomato relatives reveals orthogonal evolutionary patterns: IVs conserve underlying sequence, whereas loop anchors diversify yet recur in corresponding chromosomal neighborhoods. Finally, by integrating cross-species transcriptomics we show that promoter-anchored loops have dual regulatory roles: they associate with enhanced expression in activating chromatin contexts and with poised or repressed states when distal anchors are H3K27me3-rich. Together, this Micro-C atlas refines models of plant insulation and establishes a loop-centric, high-resolution framework to interpret plant gene regulation in 3D while providing a resource for probing how looping shapes trait variation.

## Results

### Micro-C enables high-resolution mapping of Insulation Valleys and chromatin loops in *S.lycopersicum*

To comprehensively map 3D genome structure at high resolution - including Insulation Valleys (IVs) and chromatin loops, we performed micro-C on young *S.lycopersicum* (M82) leaves and generated two independent biological replicates (see Supplementary Fig.1). These were merged into a high-resolution dataset comprising 1.45 billion informative paired-end reads, a more than 10x improvement compared to 118 million reads from the highest-resolution Hi-C dataset available for tomato^16^.

At the chromosome scale and bin sizes of 50-100kb, Hi-C and Micro-C produced highly similar ICE-balanced contact maps (Supplementary Fig. 2), but Micro-C revealed finer structural details at higher resolution (Fig. 1b, Supplementary Fig. 3). Despite minor differences, more than 98% of the genome retained consistent A/B compartmentalization between datasets (Supplementary Fig. 4).

**Figure 1.**
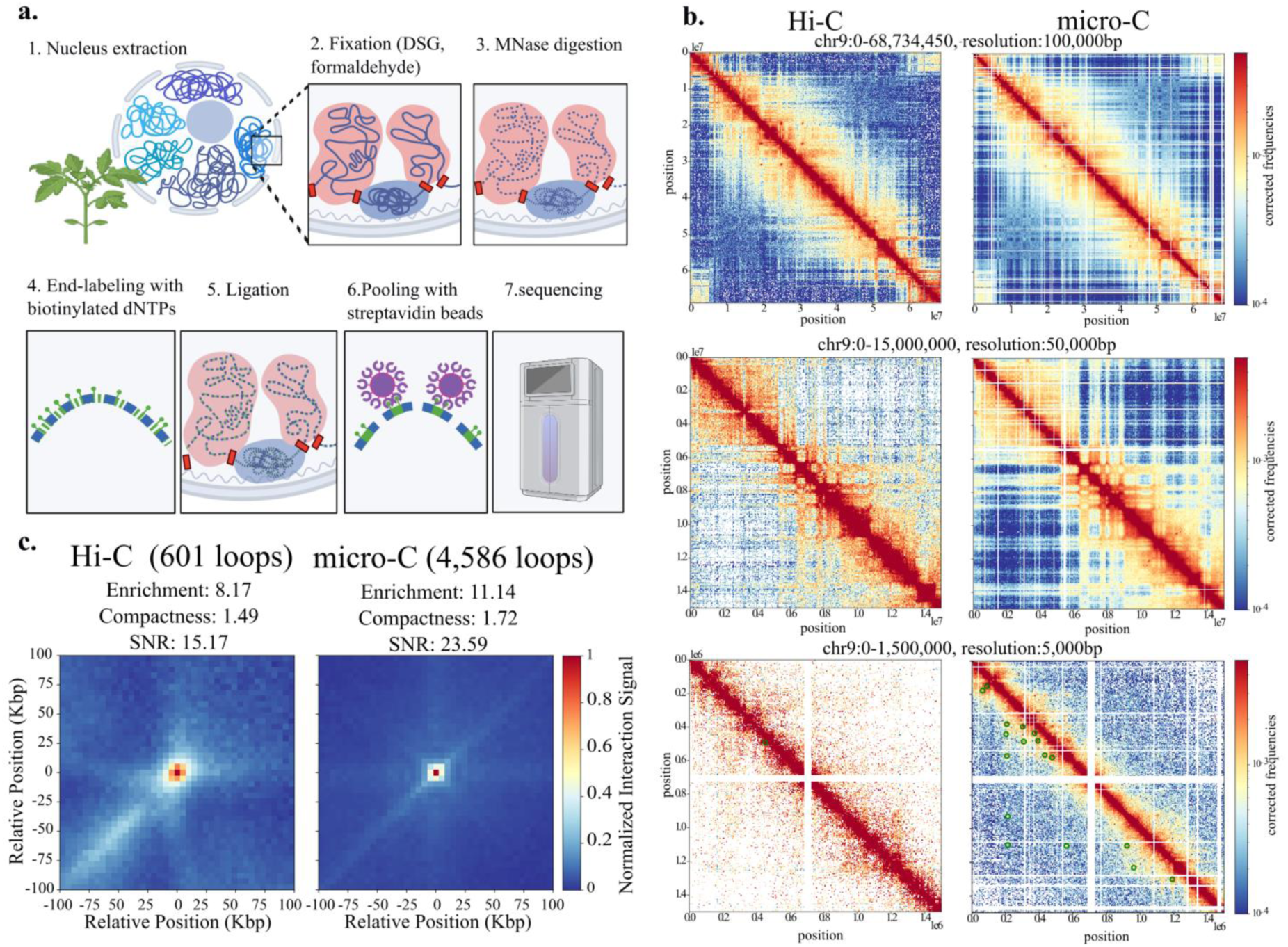
Comparison of Hi-C and Micro-C in *Solanum lycopersicum*. (a) Overview of the plant Micro-C protocol. Nuclei are extracted from tomato leaves and fixed with DSG and formaldehyde. The chromatin is then digested with MNase, and the resulting DNA ends are labeled with biotinylated dNTPs before ligation to form DNA dimers. Ligated fragments are captured using streptavidin beads and subsequently sequenced. (b Genome-wide contact maps from Hi-C and Micro-C at three resolutions. Green dots in the lower right panel mark chromatin loops called by Mustache. (c) Loop pile-ups for Hi-C (left) and Micro-C (right). Numbers above each plot report Enrichment (mean signal in a 3×3 central window divided by the mean of the entire (min–max–normalized) pileup (O/E)), Compactness (ratio of the 3×3 center mean to the mean of a surrounding 5×5 window, reflecting local sharpness.), and SNR (the maximum value in the 3×3 center divided by the standard deviation of all pixels in the pileup).

Micro-C enabled significantly improved detection of IVs by HiCExplorer^30^ and chromatin loops by Mustache^31^, yielding a fourfold increase in identified IVs (19,668 IVs detected from Micro-C data vs. 4,907 IVs from Hi-C data) and over a sevenfold increase in detected chromatin loops (4,586 loops detected from Micro-C data vs. 601 loops from Hi-C data) compared to previously published Hi-C data. Although IVs were not entirely identical between the two methods, nearly 50% of the Hi-C IVs overlapped with micro-C when allowing a ±5 kb tolerance, increasing to 100% identity at a ±25 kb threshold (Supplementary Fig. 5). Chromatin loops exhibited only 1.7% direct overlap between Hi-C and micro-C, but with a ±25 kb tolerance, 30% of Hi-C-detected loops were also found in the micro-C data (Supplementary Fig. 5). To examine whether the higher number of detected loops in Micro-C could be influenced by chromatin accessibility, we tested the bidirectional overlap between ATAC-seq peaks and loop anchors (Supplementary Fig. 6). We found that 35% of loop anchors overlap with an ATAC-seq peak, and conversely, just 9% of ATAC-seq peaks coincide with loop anchors, suggesting that increased loop detection is not primarily driven by accessibility bias. Compartment analysis further showed that IVs and loop anchors populate both A (euchromatic) and B (heterochromatic) compartments (Supplementary Fig. 7), ruling out chromatin accessibility as the primary driver of their detection.

Furthermore, the enhanced resolution of Micro-C allowed for more precise chromatin loop detection, as reflected in loop pile-up analysis (Fig. 1c). These results highlight the power of Micro-C in resolving higher-order chromatin structures with greater precision, revealing a significantly higher number of chromatin loops and IVs compared to Hi-C, while maintaining overall genome-wide compartmentalization patterns between the two datasets.

### Epigenetic Landscapes Define Chromatin Structural Features in *S.lycopersicum*

To explore the specific histone modifications that are associated with insulation valleys (IVs), we first divided IVs into promoter-associated (n = 3,805) and non-promoter (n = 4,264) groups for downstream analyses (Supplementary Fig. 7). Promoter-associated IVs are defined as those whose central bin lies within ±2 kb of a transcription start site (TSS). Using published ATAC-seq, RNA polymerase II (RNAPII) ChIP-seq, and histone-modification data of S.lycopersicum (M82) young leavs^32^, each signal was normalized to control for coverage variability and plotted over ±30 kb windows centered on each IV (Fig. 2a; Supplementary Fig. 8). In agreement with previous plant Hi-C studies^16,26,32^, both promoter-associated and non-promoter IVs exhibited strong enrichment, exceeding four standard deviations above the genome-wide average (p < 0.01, Wilcoxon signed-rank test) for ATAC-seq peaks, RNAPII, and histone marks characteristic of active promoters, gene-rich euchromatin, and Polycomb-repressed (PRC2-associated) regions, alongside depletion of constitutive heterochromatin and repetitive-element marks (Supplementary Fig. 9a). Notably, H3K4me1, formed secondary peaks approximately ±5 kb from the valley centers: in promoter-associated IVs, manifesting as two flanking peaks with a clear dip at the IV midpoint (reflecting H3K4me1’s localization in gene bodies of active genes around the promoter^33,34^), whereas in non-promoter IVs, the broader ±5 kb enrichment may indicate conserved sequence at those valleys^35^.

**Figure 2.**
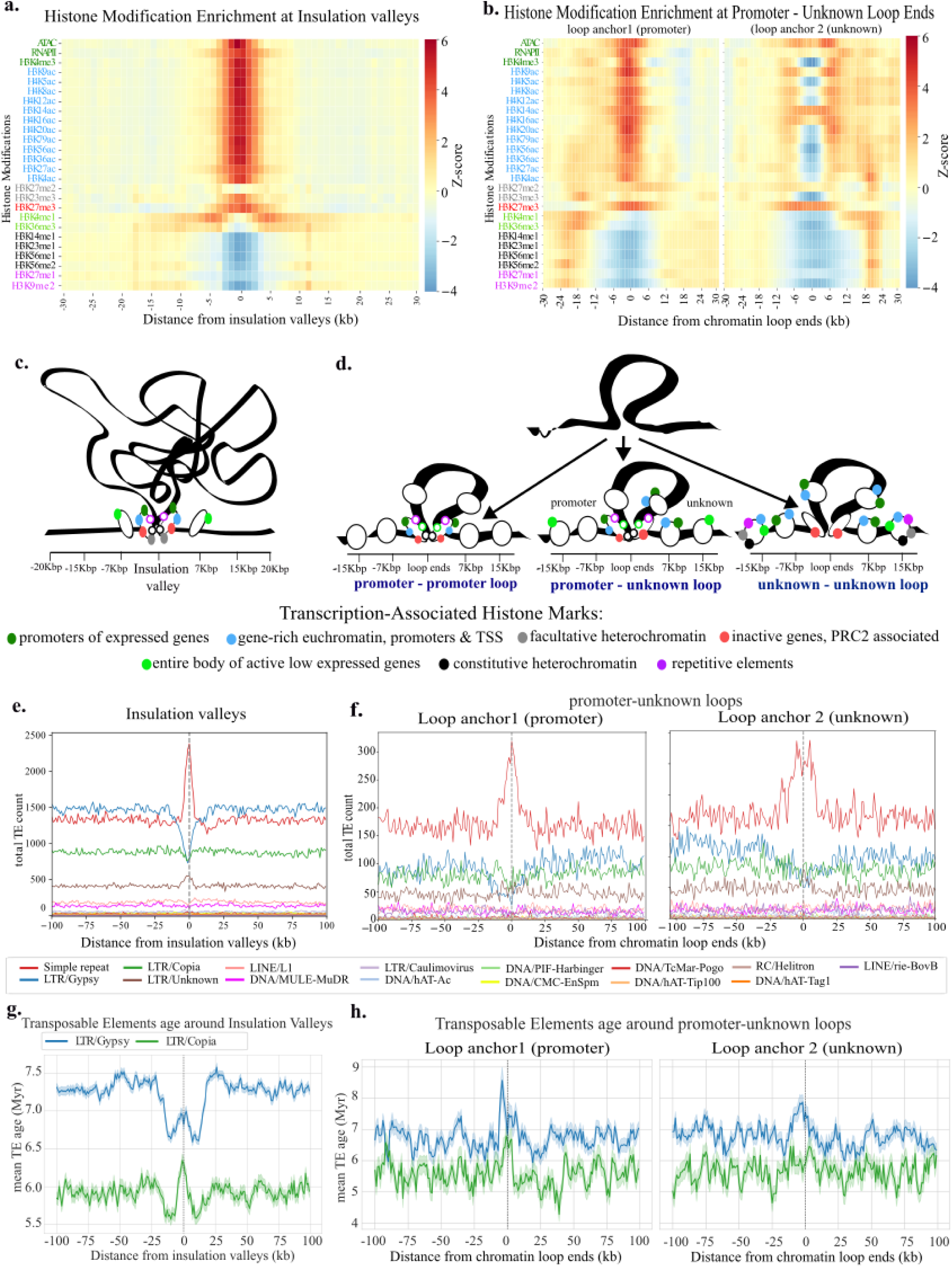
Histone modification and transposable elements at Insulation Valleys and chromatin loop anchors in *Solanum lycopersicum*. All profiles are aligned to the feature center (0; dashed line) and normalized individually to their genome-wide minimum and maximum. (a) Histone-modification heat maps (±30 kb) around IV centers, showing enrichment of promoter-like marks and depletion of heterochromatic marks at the core. (b) Histone-modification heat maps (±30 kb) around promoter–unknown (P–U) loop anchors, split by anchor identity: left, promoter; right, unknown. (c) Schematic of IV chromatin architecture: an open, RNAP II–rich core flanked by heterochromatin. (d) Schematic of loop classes: P-P (promoter-promoter), P-U (promoter-unknown) and U-U (unknown-unknown), and their characteristic chromatin environments. (e) Aggregate counts of transposable elements (TEs) by class (±100 kb) around IV centers; note enrichment of simple repeats and depletion of young LTR/Gypsy elements at the center. (f) Aggregate TE counts (±100 kb) around P-U loop anchors, shown separately for promoter (left) and unknown (right) ends. (g) Mean age of Gypsy (blue) and Copia (green) elements around IVs; shaded regions denote ± standard error of the mean. h. Mean age of Gypsy and Copia elements around P-U anchors, split by anchor identity; shading indicates ± standard error.

Next, to histone modification patterns analysis at chromatin loops, loop coordinates were aligned to the *S.lycopersicum* (M82) genome. Any loop anchor that’s laying within ±2 kb of a TSS was labeled a “promoter anchor”. Loops were then classified into four categories: promoter-promoter (P-P): 911 loops with promoters at both anchors, promoter-unknown (P-U): 778 loops with a promoter only at the first anchor, unknown-promoter (U-P): 770 loops with a promoter only at the second anchor and unknown-unknown (U-U): 2,127 loops lacking promoters at both anchors. Normalized ATAC-seq, RNAPII, and ChIP-seq signals were plotted over ±30 kb windows centered on each loop anchor (Fig. 2b, Supplementary Fig. 10).

P-P loops showed pronounced enrichment exceeding two to four standard deviations above the genome-wide average, of ATAC-seq peaks, RNAPII, and active promoter histone marks (e.g., H3K4me3, H3K9ac) at both anchors, consistent with euchromatic, actively transcribed regions (p < 0.01, Wilcoxon signed-rank test, Supplementary Fig. 10). These anchors also exhibited depletion of heterochromatin, two to four standard deviations under the genome-wide average repetitive elements, and gene-body marks associated active gene expression (p < 0.01, Wilcoxon signed-rank test). In contrast, P-U and U-P loops showed strong promoter-like signals only at their respective promoter ends, while their unknown ends were largely depleted in both active and heterochromatic marks (two to four standard deviations under the genome-wide average). However, these unknown ends exhibited a distinct H3K27me3 peak coinciding with ATAC-seq signal, along with secondary enrichment of euchromatic marks 5-10 kb from the U loop anchor. U-U loop anchors resembled the unknown ends of P-U and U-P loops, with overall lower signal but flanked (∼15-20 kb from loop anchor, outside of the loop) by peaks of constitutive and facultative heterochromatin and repetitive-element marks (four to six standard deviations above the genome-wide average, Supplementary Fig. 10).

Notably, the histone mark distribution near the anchors of U-U loops was asymmetric (Supplementary Fig. 10). For example, constitutive heterochromatin marks were more enriched ∼20kb outside the loop. Such asymmetric and directionally biased histone mark enrichment hints at a possibly directionally biases mechanisms of formation - conceptually akin to the convergent CTCF rule in vertebrates^21^, though much more work will be required to understand the mechanistic basis of this.

Insulation valleys (IVs) in *S.lycopersicum* show open-chromatin and RNAPII enrichment, whereas loop anchors show two different types of signatures: loop anchors near TSSs carry active-promoter marks; non-promoter anchors, in contrast, are marked by repressive or heterochromatic signatures flanked by euchromatic peaks. This clear separation of epigenetic states suggests that, in plants, the establishment of chromatin loops and the positioning of IVs are controlled by distinct mechanisms, hinting at a functional uncoupling of loop formation from boundary insulation.

### Depletion of active Gypsy retrotransposons around IVs and chromatin loop anchors in *S.lycopersicum*

In mammals, transposable elements such as SINEs, particularly the MIR family, and DNA repeats have been found to be enriched at CTCF binding sites, especially at TAD boundaries conserved across primates^36^. These elements have therefore been proposed as contributors to the evolution of 3D genome organization. A comparable pattern is seen in cotton, where Hi-C maps at 50 kb resolution show enrichment of Gypsy retrotransposons at TAD-like boundaries; younger Gypsy copies mark species-specific boundaries, whereas older copies mark conserved ones^37^. By contrast, high-resolution (5 kb) Micro-C in rice revealed a marked depletion of Gypsy elements at TAD boundaries, strongest at evolutionarily conserved boundaries^38^.

Analysis of transposable element (TE) distribution at *S.lycopersicum* IVs identified from Micro-C data, revealed that IVs enriched with simple repeats, such as (AT)n, (TA)n, and (T)n, and depleted of Gypsy and Copia retrotransposons, two of the dominant TE families in the tomato genome (Fig. 2e, Supp. Fig. 11). Promoter-associated IVs show depletion of both Gypsy and Copia elements, whereas non-promoter IVs are depleted only for Gypsy (Supp. Fig. 9b), suggesting that Copia depletion at promoter-associated IVs may reflect local gene density rather than IV structure per se, while Gypsy depletion is not affected by gene density. Interestingly, in promoter-associated IVs, both Gypsy and Copia retrotransposons exhibit younger mean ages approximately ±10 kb from IV centers but older mean ages precisely at the IV midpoints (Fig. 2g). When stratified by IV class, promoter-associated IVs harbor relatively older Gypsy and Copia elements throughout the ±10 kb window, while non-promoter IVs show younger Gypsy and Copia ages, particularly in the ±10 kb flanks (Supp. Fig. 9c). These results indicate that TE insertions are more tolerated around non-promoter IVs but that their centers remain relatively protected (as reflected by a slight age “valley” at the midpoint), whereas promoter-associated IVs are highly protected from new insertions.

Loop anchors similarly exhibit enrichment of simple repeats such as (AT)n and (TA)n (n_median_=40bp), particularly at promoter-associated ends, accompanied by mild depletion of Gypsy and Copia retrotransposons (Fig. 2f, Supplementary Figs. 12-13). Further analysis of transposable element (TE) age revealed distinct patterns across loop types and anchor positions (Fig. 2h, Supplementary Fig. 14). Gypsy elements exhibited a pronounced peak in mean age centered at loop anchors, especially at both anchors of P-P loops, with similar but slightly reduced peaks observed at the promoter-associated ends of P-U and U-P loops. U-U loops also showed elevated Gypsy element age at both anchors, albeit to a lesser extent. In contrast, Copia elements displayed consistently younger average ages and relatively flatter profiles, with no significant peaks near loop anchors. These results suggest that older Gypsy elements are preferentially retained or enriched near chromatin loop anchors, not only those associated with promoters but also at non-promoter (U-U) anchors, potentially reflecting a protective mechanism that limits new Gypsy insertions in these structurally constrained regions of the genome.

In summary, *S.lycopersicum* reconciles the cotton-versus-rice paradox by splitting it across chromatin features. All IVs are Gypsy-depleted, echoing the rice pattern; yet their promoter-associated subset harbors older Gypsy copies (cotton-like), whereas non-promoter IVs contain the youngest insertions. Loop anchors introduce a novel layer: they show only a mild Gypsy depletion but consistently retain ancient Gypsy elements at every anchor, irrespective of promoter status. Thus, tomato chromatin deploys feature-specific age filters, strict exclusion in IV cores and age-selective retention at loop anchors, to manage Gypsy activity.

### Convergent structure, divergent mechanism: plants match mammalian IV/loop scales, but only mammals show an extrusion shoulder

Because plants lack CTCF, the architectural anchor of mammalian structural loops and TADs, we compared loop- and IV-based 3D genome organization between *S.lycopersicum* and mammals to understand conserved and divergent principles. Micro-C has already charted high-resolution chromatin architecture in mouse^24^ and human embryonic stem cells^39^, and in *A.thaliana*^26^. extending this view to tomato lets us test whether insulation valleys (IVs) and loops follow similar length scales, epigenetic landscape and IV-loop coupling in plants.

To eliminate sequencing-depth bias, we down-sampled the tomato Micro-C data to the same read coverage as the mESC Micro-C^24^, before calling IVs and loops. The mESC normalized map contains 17,886 IVs (mean spacing = 143 kb; N50 = 210 kb), whereas tomato has 7,210 IVs (mean = 111 kb; N50 = 150 kb) (Fig. 3a, left; Supp. Fig. 15). Although the spacing distributions differ statistically (Mann-Whitney U = 47,497, *p* < 0.001; KS D = 0.126, *p* < 0.0001), both fall within the ∼100-200 kb window reported across plants and mammals. Loop sizes echo the same pattern; mESCs harbour 23,532 loops (mean = 515 kb; N50 = 835 kb), while tomato shows 4,261 loops (mean = 496 kb; N50 = 830 kb) (Fig. 3a, right; Supp. Fig. 15). The distributions again differ (Mann-Whitney U = 53,208,083, *p* < 0.0001; KS = 0.0548, *p* < 0.0001), yet both cluster around the conserved ∼300-400 kb size class.

**Figure 3.**
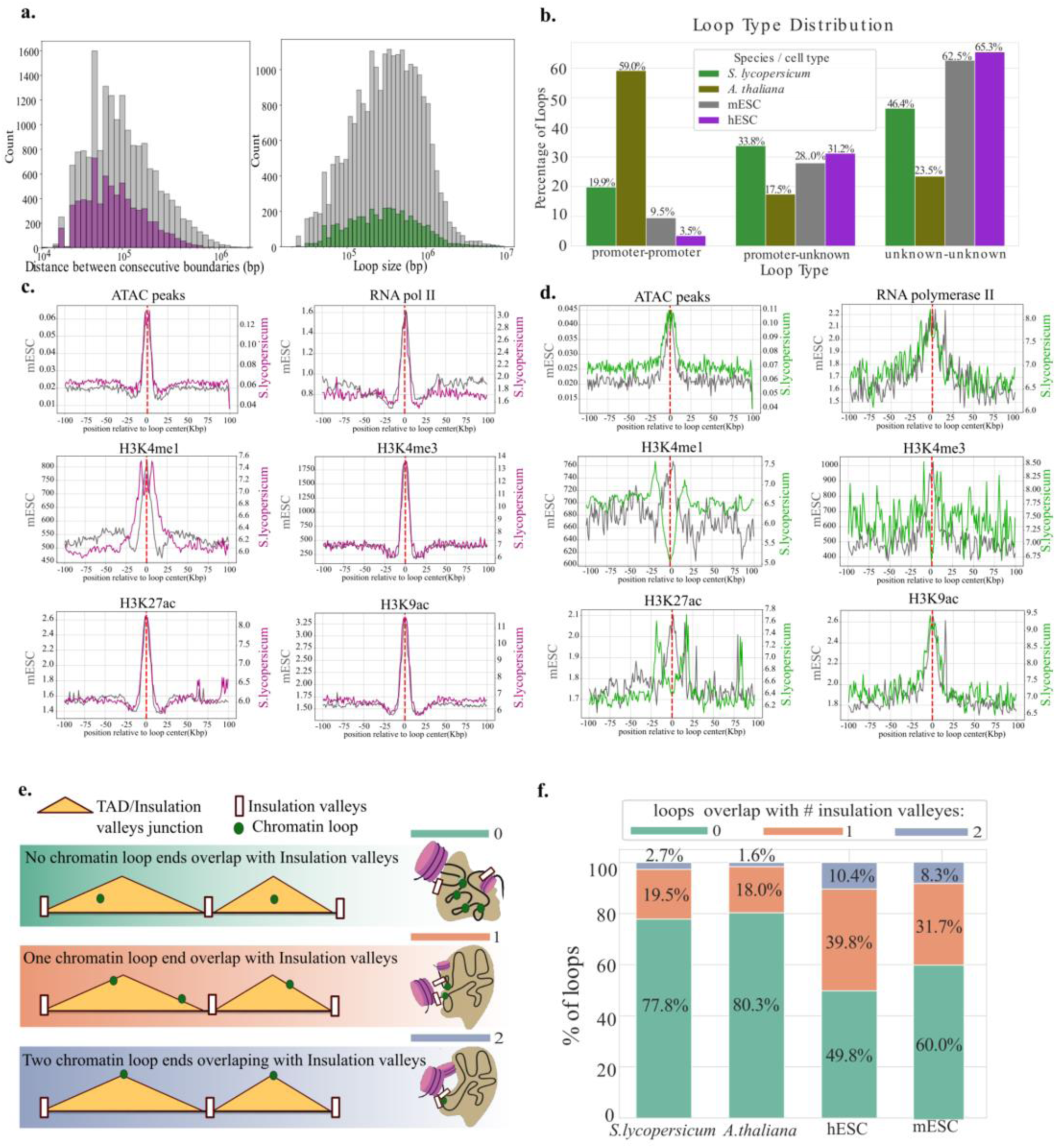
TAD-like Structures and Chromatin Loops across plant and mammalian genomes. (a) Left: distribution of distances adjacent insulation-valley midpoints in *S.* lycopersicum (purple) and mESCs (grey). Right: histogram of loop-span lengths (bp) in *S.lycopersicum* (green) and mESCs (grey), on a log₁₀ scale. (b) Stacked bars showing the percentage of loops classified as Promoter-Promoter, Promoter-Unknown or Unknown-Unknown in *S.lycopersicum*, *A.thaliana*, human ESCs (hESC) and mouse ESCs (mESC). (c) Aggregate enrichment profiles of ATAC-seq, RNA polymerase II, H3K4me1, H3K4me3, H3K27ac, and H3K9ac signals around insulation sites in *S.lycopersicum* (purple) and mESCs (grey). (d) Equivalent aggregate profiles around chromatin loop anchors in *S.lycopersicum* (green) and mESC (grey). (e) Schematic illustrating loop-anchor overlap with insulation valleys: 0 = no overlap, 1 = one end overlaps, 2 = both ends overlap. (f) Frequency of loop-valley overlap categories (0, 1 or 2) for each species/cell type; bar segments colored as in (e).

A simple polymer view predicts that densely packed, unknotted chromatin (a “fractal globule”) shows contact probability that scales roughly as *P(s)* ∝ *s*⁻¹ over mid-range genomic distances; this *s⁻¹* behavior was observed in early Hi-C data and formalized in polymer theory^40,41^. Consistently, depth-normalized 1-kb *P(s)* profiles are broadly similar across plants and mammals, following an ≈*s⁻¹* decay from ∼10⁴-10⁶ bp, indicating shared baseline polymeric folding (Supp. Fig. 15). A clear shoulder at 10⁵-10⁶ bp, with the derivative rising toward 0, is observed only in hESC/mESC, consistent with CTCF-stalled cohesin extrusion; plant profiles (three *Solanum* species and *Arabidopsis*) lack this feature and show a smooth decay with a more negative, monotonic slope. These analyses therefore suggest that while global scaling is conserved, extrusion in plants is absent, weaker, or operates without a barrier-defined characteristic scale. Thus, despite the absence of CTCF in plants, the characteristic length scales of IVs and loops are remarkably similar in tomato and mammals, whereas stabilized point loops are a mammalian hallmark.

### *S.lycopersicum* Loop Anchors Have Distinct Epigenetic Signatures and Rarely Coincide with Insulation Valleys, in Contrast to mESCs

To broaden the scope, we extended the promoter-loop anchor proximity analysis to mammalian genomes and a wider diversity of plant species, classifying an anchor as promoter-associated when its midpoint lay within ±2 kb of a TSS. In plants, a larger fraction of loops are promoter-promoter (P-P) compared with mammals, the promoter-unknown (P-U) fraction is similar in both, and the unknown-unknown (U-U) group is smaller in plants (Fig. 3b; Supp. Fig. 16).

Comparison of histone-modification patterns at IVs (±100 kb) between mESCs and *S.lycopersicum* revealed remarkably similar enrichments of ATAC-seq peaks and RNA-polymerase II ChIP-seq signals, like the active-promoter marks H3K4me3, H3K27ac and H3K9ac in both mammalian and plant promoters (Fig 3c). By contrast, H3K4me1which is-associated with mammalian active enhancers^42,43^ and with gene bodies of actively expressed plant genes^33,34^, showed a single sharp peak at insulation valley centers in mESCs but three distinct peaks in *S.lycopersicum,* reflecting its divergent role in the two lineages. A similar comparison at chromatin loop anchors showed that ATAC peaks, RNA-polymerase II, and H3K9ac patterns are broadly conserved between *S.lycopersicum* and mESCs; however, *S.lycopersicum* loop anchors display an inverted H3K4me1 profile, altered H3K27ac enrichment, and essentially no H3K4me3 signal (Fig. 3d). In mESCs, all six marks (ATAC, RNAPII, H3K4me1, H3K4me3, H3K27ac, H3K9ac) form crisp, single peaks at both IVs and loop anchors, whereas *S.lycopersicum* loop anchors lack those peaks for H3K4me1, H3K4me3, and H3K27ac, implying that plant loop anchors carry distinct epigenetic signatures and that, in tomato, loops and IVs rarely coincide.

To gain further insight into the relationship between IVs and chromatin loops, we defined three overlap scenarios (Fig. 3e): Zero overlap: neither loop anchor overlaps an IV, Single overlap: one loop anchor overlaps an IV or Double overlap: both loop anchors overlap IVs. When we quantified these categories in both plant and mammalian cells, we observed a clear difference in overlap frequency (Fig. 3e; Supp. Fig. 17). In mammals, 8.3-16.9 % of loops have both anchors overlapping IVs, compared with only 0.5-3.1 % in plants. Likewise, 31.7-45.4 % of mammalian loops have exactly one anchor overlapping an IV, whereas just 7.3-22.6 % of plant loops fall into that category. Hence, unlike the tight, TAD boundary-centric looping seen in mammals, plant genomes appear to decouple insulation and looping: the vast majority of tomato loops anchor outside IV cores, revealing a plant-specific, boundary-independent mechanism for loop formation.

### Insulation Valleys Are Sequence-Conserved Whereas Loops Retain Positional Context Across *Solanum* Species

In mammals, studies have shown that 3D genomic features are conserved across diverse species, underscoring their fundamental role in maintaining gene regulation and genome integrity^36^. To investigate whether similar evolutionary conservation exists in plants, we performed Micro-C in two wild relatives of *S.lycopersicum*: *S.pimpinellifolium* (LA1589), which diverged approximately 1 million years ago, and *S.pennellii* (LA0716), which diverged around 5 million years ago (Fig. 4b).

**Figure 4.**
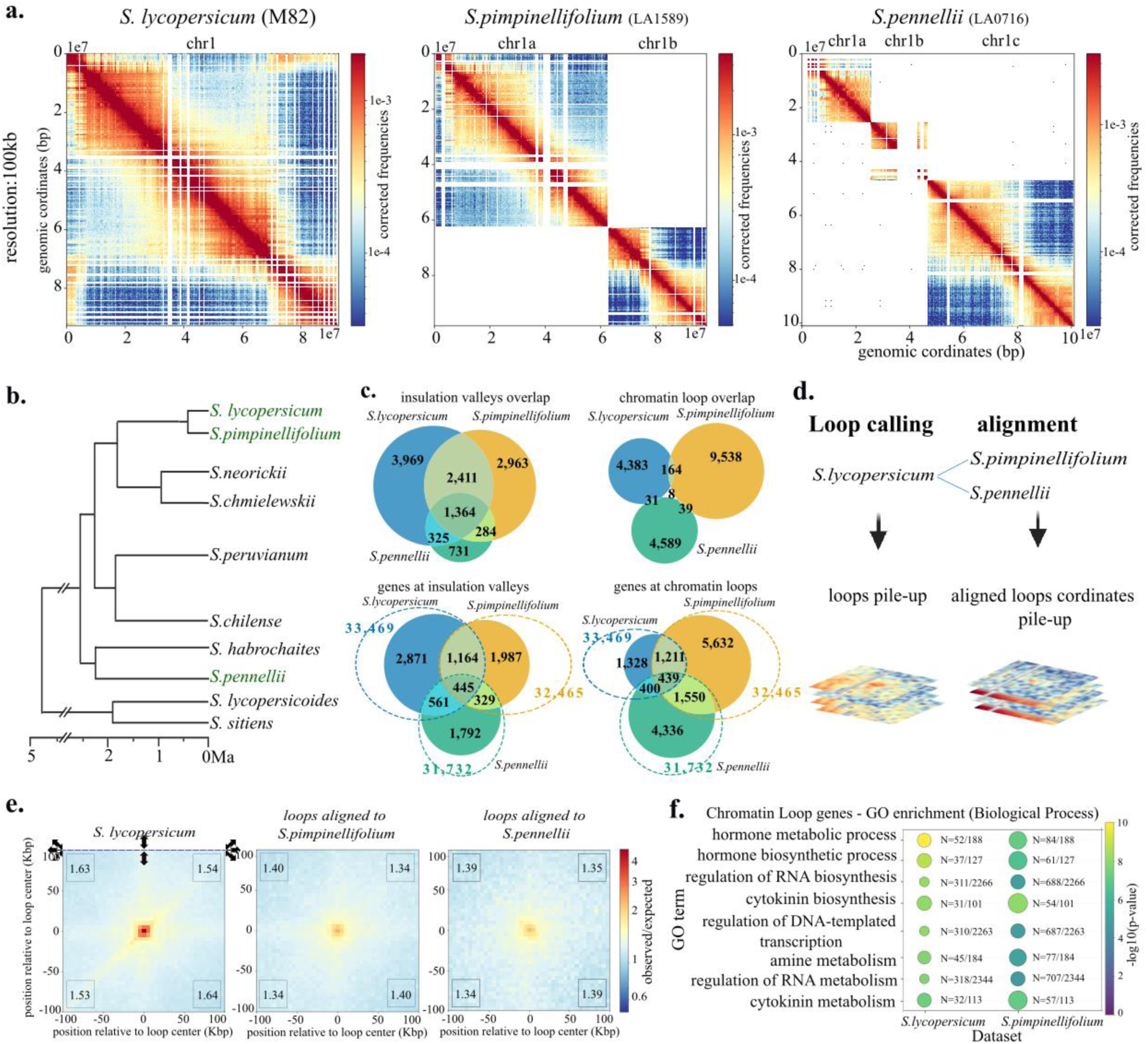
Conservation of Insulation Valleys and Chromatin Loops in Solanaceae. (a) Whole chromosome contact heatmaps for *S.lycopersicum* compared to *S.pimpinellifolium* and *S.pennellii* (resolution: 100 kb). (b) Phylogenetic tree of the Solanaceae family, with the three species analyzed in this study highlighted in green (based on Pease et al.^67^). (c) Top panel: *(Left)* Number of shared insulation valleys between *S.lycopersicum* and each relative. *(Right)* Number of shared chromatin loops based on sequence conservation. Bottom panel: (*Left)* Genes whose transcription start sites (±2 kb) lie in insulation valleys and are present in *S.lycopersicum* plus one wild relative. *(Right)* Genes whose transcription start sites (±2 kb) coincide with chromatin-loop anchors and are shared between *S.lycopersicum* and the indicated relative. The numbers in dashed spheres represents the total number of homolog genes in each genome (d) Schematic of the aligned pile-up analysis. Loops identified in *S.lycopersicum* were lifted over to *S.pimpinellifolium* and *S.pennellii*, and aligned anchors were aggregated. The numbers at the corners of each pile-up plot indicate the ratio between corner signal and center signal. (e) Pile-up plots: Left - self- pileup of *S.lycopersicum* loops; Middle - *S.lycopersicum* loops lifted over to *S.pimpinellifolium*, *S.pimpinellifolium* aligned coordinate are piled-up; Right - *S.lycopersicum* loops lifted over to *S.pennellii*. *S.pennellii* aligned coordinate are piled-up; (f) GO-term enrichment (biological-process category) for genes located at conserved chromatin-loop anchors in *S.lycopersicum* and *S.pimpinellifolium*.

To minimize artifacts caused by genomic rearrangements, commonly observed in wild tomato species (see Supplementary Fig. 18), we sequenced both accessions using PacBio long-read technology and generated *de novo* reference assemblies. The resulting genomes contained 26 contigs >1 Mb for *S.pimpinellifolium* and 75 contigs >1 Mb for *S.pennellii* (Supplementary Fig. 18). Then, Micro-C contact maps from all three species were aligned, revealing broadly similar interaction patterns (Fig. 4a). At the chromosome level, *S.lycopersicum* and *S.pimpinellifolium* exhibited strong conservation of chromatin structure (Pearson > 0.96, SSIM ∼0.99–0.995), while *S.pennellii* showed moderate to high conservation. Several chromosomes in the *S.lycopersicum* vs. *S.pennellii* comparison had Pearson values below 0.7, suggesting the presence of genomic rearrangements or structural divergence (Supplementary Fig. 19).

To assess the sequence conservation of specific structural features, we analyzed insulation valleys (IVs) and chromatin loops across the three species. Using 5-kb bins, we aligned IV regions from each species to the other genomes and defined shared IVs as those whose aligned regions also overlapped IVs in the target species. We found that ∼43% of *S.lycopersicum*, 63% of *S.pimpinellifolium*, and 94% of *S.pennellii* IVs were conserved (Fig. 4c, top panel, left), indicating substantial sequence conservation. IVs were more frequently shared between *S.lycopersicum* and *S.pimpinellifolium* than between *S.lycopersicum* and *S.pennellii*, suggesting that structural conservation correlates with evolutionary distance. Moreover, as reported for mammals^36^, cotton^37^, and rice^38^, the evolutionarily conserved tomato IVs display lower insulation scores than species-unique IVs (Supplementary Fig. 21), consistent with stronger or more stable boundaries.

Unlike the well-conserved nature of IVs, chromatin loop anchor sequences were poorly conserved. Fewer than 1% of loop anchor sequences were shared among all three species, and only 1-5% were shared between any two species (Fig. 4c, top panel, right). Nonetheless, conserved loop anchor sequences have higher contact frequency or prevalence across cells as implied by the significantly lower FDRs (Supplementary Fig. 21). To test whether slight anchor position shifts mask deeper conservation, we aligned each *S.lycopersicum* loop anchor sequence (5-kb) to the other genomes and generated contact pile-ups at the aligned coordinates (Fig. 4d,e). Despite the rapid sequence turnover at anchor sites, strong contact enrichment recurs at the same chromosomal neighborhoods in all three species. Thus, while IVs conserve their underlying DNA sequence, loops conserve their positional context along the chromosome, even as the anchor sequences themselves diverge.

### Chromatin Loops Anchor Evolutionarily Stable Genes Involved in Key Regulatory Processes

To explore the functional conservation of chromatin structures, we next asked whether the genes near IVs and loop anchors are conserved across species. We lifted over the *S.lycopersicum* genome annotation to *S.pimpinellifolium* and *S.pennellii*, then identified genes with transcription start sites (TSSs) located within ±2 kb of either IV centers or loop anchors. Comparing gene identities (Fig. 4c, bottom panels) revealed that a similar number of genes are shared at loop anchors and IVs, suggesting comparable evolutionary constraints.

Gene Ontology (GO) enrichment analysis of genes near loop anchors showed significant enrichment for essential biological processes in both *S.lycopersicum* and *S.pimpinellifolium* (Fig. 4f, Supplementary Figs. 22-23). Enriched terms included hormone biosynthesis and signaling, particularly cytokinin, as well as nucleic acid metabolic and biosynthetic processes. These findings are consistent with the developmental context of the samples (young leaves undergoing active growth and division) and suggest that chromatin loops preferentially engage regulatory regions of genes involved in fundamental cellular functions.

In conclusion, our results show that while chromatin loops are less conserved at the sequence level than IVs, they preferentially anchor conserved regulatory genes and maintain their spatial positioning. This highlights their likely importance in sustaining core regulatory programs during evolution.

### Promoter-anchored chromatin loops have dual roles in enhancing and poising gene expression

To characterize how chromatin loops relate to transcription across *Solanum*, MARS-seq^44^ was performed on RNA from young leaves of ten biological replicates per species (*S. lycopersicum*, *S. pimpinellifolium*, *S. pennellii*). Duplicate cDNA molecules were removed with the UTAP pipeline^45^. Genes whose promoters lie at chromatin-loop anchors (Fig. 4c, bottom left) were first filtered for expression (mean TP100k > 0.05 in all species). or each gene, a cross-species loop-expression contrast was computed as Δ=mean[log_2_(TP100k)]_looped_ -mean[log_2_(TP100k)]_non-looped_ (figure 5a), where means are taken across species in which gene is looped or non-looped, respectively. The resulting distribution is continuous, spanning strongly positive (Δ > 8) to strongly negative (Δ < -5) values, indicating that promoter-anchored loops can be associated with both enhanced and repressed/poised expression.

**Figure 5.**
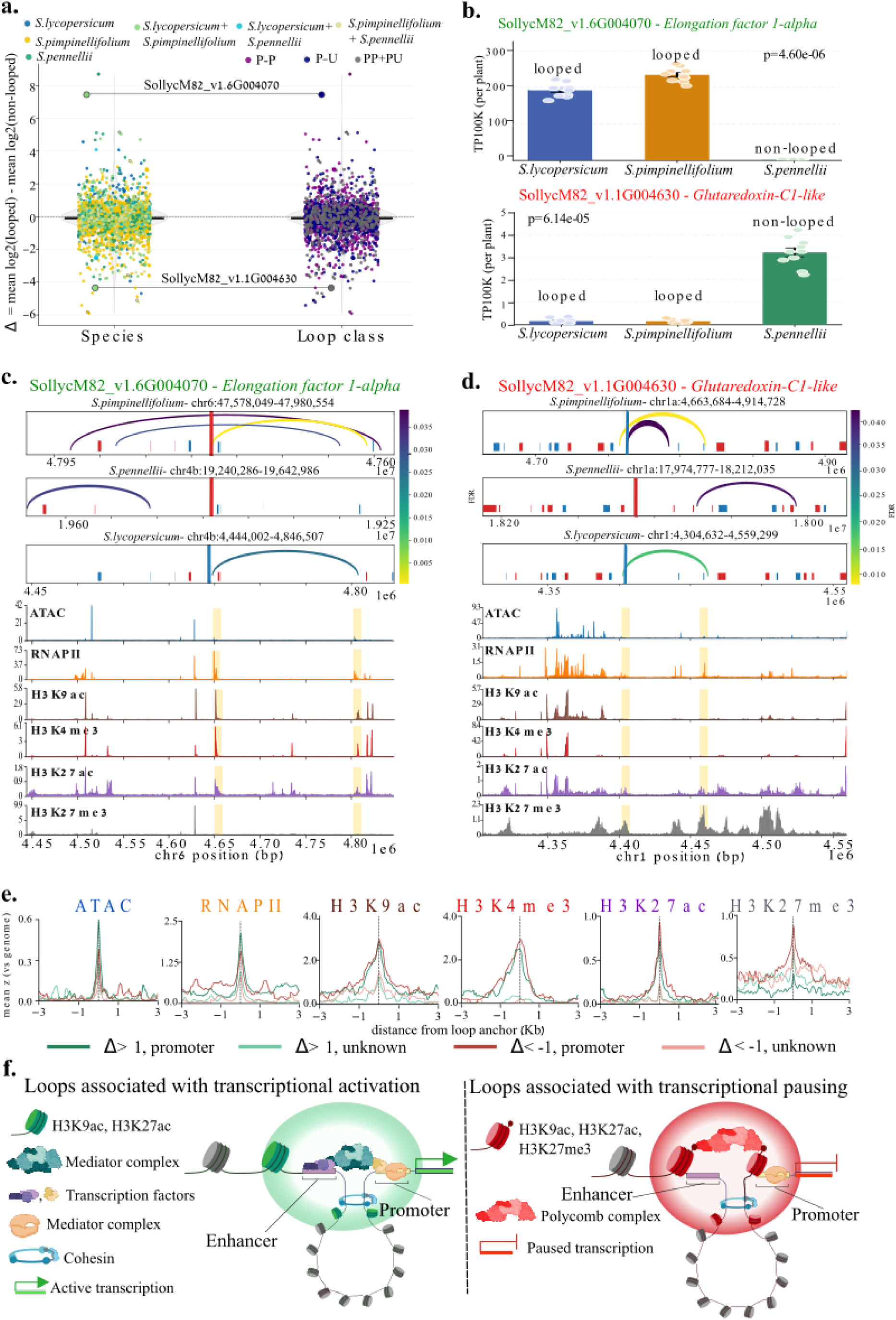
Chromatin looping shows graded coupling to transcriptional activation and pausing across tomato species. (a) Looped vs non-looped genes show a continuum of expression differences, consistent with activating, repressing, and neutral loop contexts. For each expressed anchor gene present in *S. lycopersicum*, *S. pimpinellifolium*, and *S. pennellii* (mean TP100k > 0.05 in all three), Δ=mean[log_2_(TP100k)]_looped_-mean[log_2_(TP100k)]_non-looped_ is plotted. Each dot represents one gene. Left panel: points colored by the species in which the gene is looped. Right panel: points colored by loop class at the promoter-P-P (promoter-promoter), P-U (promoter-unknown), or PP+PU (the gene occurs at multiple loops, including P-P and P-U). The Δ distribution is continuous, indicating dual, and graded, roles for looping: some loops correlate with enhanced expression, others with repressed/poised expression, and a subset show little or no association with expression. (b) Examples. Top: *Elongation factor 1-alpha* (SollycM82_v1.6G004070) shows a positive Δ (looped states associated with higher expression). Bottom: *Glutaredoxin-C1-like* (SollycM82_v1.1G004630) shows a negative Δ (looped states associated with reduced/poised expression). Bars show mean TP100k per species; points denote biological replicates; *p*-values (Kruskal-Wallis, one-way ANOVA) compare looped versus non-looped states. (c-d) Genome views for the two genes in (b). Locus-level track panels for the two genes in (b). Upper tracks: chromatin-loops in each species; arc color encodes FDR. Forward-strand genes are blue; reverse-strand genes are red; the focal gene is emphasized with thicker gene track. Lower tracks: *S. lycopersicum* chromatin profiles (ATAC-seq, RNAPII, H3K9ac, H3K4me3, H3K27ac, H3K27me3). Loop anchors are highlighted with a yellow background. Panel c: *Elongation factor 1-alpha*; panel d: *Glutaredoxin-C1-like*. (e) Peak-centered metaprofiles (±3 kb; mean z-score) of *S. lycopersicum* ATAC-seq, RNAPII, H3K9ac, H3K4me3, H3K27ac, and H3K27me3 at P-U loops. Curves are stratified by gene loop-expression associetion: Δ > 1 (loops associated with enhanced expression, colored in greens) versus Δ < -1 (loops associated with repressed/poised expression, colored in reds). Promoter anchor signals are in vivid colors and unknown anchors in lighter colors. The vertical dashed line marks the anchor center. (f) Working model. Left: loops associated with transcriptional activation; transcription factors are bound to the non-promoter anchor, which enriched for H3K9ac/H3K27ac and accessible chromatin; Mediator and RNAPII are recruited to the promoter, leading to increased transcription. Right: loops associated with transcriptional pausing; both anchors bear high H3K27me3; Polycomb complexes bind methylated histone tails and restrain RNAPII, yielding a paused/repressed state. *Abbreviations:* TP100k, transcripts per 100,000.

To illustrate these trends, genes with extreme Δ values were examined (Fig. 5b; Supplementary Fig. 24). At the *Elongation factor 1-alpha* gene, gene with positive Δ, the promoter and partner anchors display strong ATAC-seq, RNAPII, and activating histone marks (H3K9ac, H3K4me3, H3K27ac) at both anchors of P-U loops (Fig. 5c). Conversely, at the *Glutaredoxin-C1-like*, gene with negative Δ, there is a high H3K27me3 at both anchors with low RNAPII signal and no H3K9ac, H3K4me3 signals (Fig. 5d).

To determine whether these patterns extend genome-wide rather than reflecting a few exemplars, histone-mark enrichment was quantified at *S.lycopersicum* promoter-unknown (P-U) loops stratified by Δ, where Δ>1 denotes ≥2-fold higher expression in looped versus non-looped genes and Δ<-1 denotes ≥2-fold lower expression. We focused on P-U loops to minimize promoter-associated signal at both anchors and to isolate the behavior of the distal, non-promoter anchor. Promoter anchors showed similar mean peak levels of RNA polymerase II, H3K9ac, and H3K27ac in loops associated with enhancement (Δ>1) and repression (Δ<−1), whereas the unknown anchors displayed smaller peaks for these activating marks (Fig. 5e; Supplementary Fig. 24). In contrast, both the promoter and the unknown anchors of repressed loops (Δ<-1) exhibited higher H3K27me3 compared with enhanced loops (Δ>1).

Taken together, these results support a dual-role model for looping (Fig. 5f). In an activating configuration, the distal anchor recruits transcription factors and cofactors (e.g., Mediator), creating a permissive environment that increases RNAPII occupancy and transcription; non-looped genes therefore show relatively lower expression. In a poising configuration, the distal anchor is also enriched for H3K27me3 and engages Polycomb Repressive Complex 2 (PRC2), promoting H3K27me3 spreading and dampening transcription. In this case RNAPII at the promoter is not markedly reduced, but transcription is repressed; non-looped genes exhibit relatively higher expression because the distal environment is less repressive.

## Discussion

Delivering a nucleosome-scale contact map, our tomato Micro-C dataset establishes a new benchmark for plant 3-D genomics and uncovers an architectural logic that departs sharply from the CTCF-centric paradigm in animals, while resolving several ambiguities left by earlier plant Hi-C studies.

Consistent with promoter-capture Hi-C studies in Arabidopsis^46^ and maize^47^, our promoter-agnostic maps, shows that more than half of tomato loops link a promoter to either another promoter or a distal open site. Yet, unlike mammalian CTCF/cohesion-anchored loops, tomato loops rarely originate from insulation valleys (IVs). This uncoupling points to distinct formation routes: IVs likely arise from sequence or chromatin features that keep loci open and RNAP II-rich, whereas loops assemble wherever transcriptional or Polycomb complexes converge. One possibility is that, without an obligatory motif or orientation, loops nucleate wherever compatible bridging complexes accumulate-transcriptional machinery at promoters or Polycomb condensates at repressed loci. In this motif-free scenario, P-U pairs naturally yield “mixed” chromatin signatures: the promoter side is euchromatic, whereas the distal partner can be Polycomb-enriched or euchromatic, so averaging anchors produces a composite profile. Alternatively, plants may harbor oriented sequence or protein tethers that position loops independently of IVs. Such elements would be invisible to boundary-centric analyses yet could still enforce directionality, as hinted by U-U loops, where we observe asymmetric heterochromatic enrichment ∼20 kb outside of the loop in assymetric pattern. This pattern suggests specific, possibly oriented features that stabilize loops without coinciding with insulation architecture.

Enhancers underpin long-range regulation in animals; however, their plant counterparts have been hard to characterize. Only a handful of bona-fide examples are known (for review^48^), canonical marks are less predictive and bidirectional eRNAs are rare^23^. Distal anchors that recur at syntenic positions across multiple *Solanum* species emerge as especially strong enhancer candidates. For example, the GAME enhancer 1 (GE1)^49^ region forms a strong loop hub in *S.pimpinellifolium* (LA1589) carrying an activating GE1 allele, but not in *S. lycopersicum* (M82) or *S. pennellii* (LA0716) (Supp. Fig. 24), spotlighting a tractable locus for CRISPR interference, reporter assays, and allele-swap tests near metabolic, hormone, and development genes enriched among looped loci.

Evolutionary comparison sharpens the contrast between IVs and loops. Solanum IVs were shown to be sequence-conserved across tomato, potato, eggplant, and pepper, underscoring strong purifying pressure on domain borders^50^. Our comparative Micro-C maps extend this view and reveal that IVs and loops obey distinct evolutionary logics. IVs remain largely sequence-locked: 40–60 % of boundaries in cultivated tomato align with those in two wild relatives and stay almost free of young Gypsy insertions. A nuance emerges when IVs are stratified by gene context: promoter-associated IVs exclude young Gypsies altogether, whereas non-promoter IVs tolerate a modest influx of recent copies, hinting at a hierarchy in which non-promoter boundaries are more amenable to evolutionary tweaking.

Loops anchors show little primary-sequence conservation and milder Gypsy depletion, yet they repeatedly tether the same suites of orthologous promoters involved in hormone signaling, RNA metabolism and development. The resulting “loop paradox” can be explained if the DNA sequence cues that nucleate plant loops are short, mobile and functionally redundant: local motif drift, small indels, or transposon insertions can shuffle these elements within a permissive cis-window without disrupting the 3-D contact. Selection thus acts on maintaining spatial proximity rather than nucleotide identity.

Across plant studies, loop-expression relationships have pointed in opposite directions depending on assay bias: open-chromatin-anchored maps tend to recover activating contacts, whereas conventional Hi-C often highlights Polycomb-linked, repressive long-range contacts. Our unbiased Micro-C resolves both within a single framework and indicates that the distal anchor’s chromatin state, rather than loop size or PP/PU class, best predicts transcriptional outcome (Supplementary figure 26). Unlike cotton, where very long or highly networked loops were associated with lower expression^29^, we find no association between loop length, loop type, loop species and Δ (enhanced vs repressed expression). Key open questions now become tractable with the tomato Micro-C atlas: (i) do short, distal sequence elements suffice to nucleate loops, and are they orientation-independent? (ii) are loops functionally monovalent (intrinsically enhancing or repressing) or bivalent, switching outcome with the histone environment at either anchor? Addressing these will require anchor fine-mapping and perturbation, CRISPR deletions/swaps of distal elements, chromatin editing to flip H3K27me3/acetylation at anchors, and targeted disruption of Mediator/PRC2, plus multiplex contact assays to test whether additional partners modulate loop polarity.

Together, our data support a model in which sequence-encoded IVs form stable chromatin “walls,” while activity-driven loops act as adjustable “wires” across them. Crucially, promoter-anchored loops can either enhance or repress transcription depending on the distal anchor’s chromatin state, boosting expression when paired with accessible/acetylated chromatin, but poising or repressing when H3K27me3/PRC2-enriched. Dissecting the proteins, RNA molecules and short DNA sequence or epigenetic cues that place anchors, testing enhancer activity at non-promoter sites, and perturbing histone states to flip loop outcomes now become tractable goals, with direct relevance for tuning traits in crop improvement.

## Methods

### Plant materials and growth conditions

*Solanum lycopersicum* cv. M82, *S. pimpinellifolium* LA1589 and *S. pennellii* LA0716 seeds were germinated on Nitsch medium (0.2 % MS, 2 % sucrose, 1 % agar, pH 5.8) and grown at 25 °C, 70 % relative humidity under a 16 h light / 8 h dark photoperiod. Young leaves were harvested from 3-week-old seedlings.

### Micro-C library preparation and sequencing

For each genotype, leaves from ∼60 seedlings were pooled. Nuclei were released by chopping tissue in ice-cold LB01 buffer (15 mM Tris-HCl pH 7.5, 2 mM EDTA, 0.5 mM spermine, 80 mM KCl, 20 mM NaCl, 0.1 % Triton X-100) for 15 min and incubated on ice for 10 min. After three washes (450 ×g, 4 °C, 30 min each) in cold BSA buffer, nuclei were processed exactly as in the mammalian Micro-C protocol of Goel et al.^51^. Libraries were sequenced on an Illumina NovaSeq X platform at the Nancy and Stephen Grand Israel National Center for Personalized Medicine (G-INCPM), Weizmann Institute of Science.

### Whole-genome sequencing and *de-novo* assembly

High-molecular-weight genomic DNA from *S.pimpinellifolium* LA1589 and *S.pennellii* LA0716 was extracted using CTAB^52^ and sequenced with the PacBio® Revio platform (Maryland Genomics, University of Maryland, USA). HiFi reads were *de-novo* assembled using hifiasm^53^. Contigs were aligned to the M82 reference with Minimap2^54^, and renamed according to their best syntenic M82 chromosome to facilitate cross-species comparisons.

### Micro-C data processing

Reads were aligned to the M82 genome^55^ or the newly assembled *S.pimpinellifolium* LA1589 and *S.pennellii* LA0716 genomes using Bowtie2^56^ ( --reorder --very-sensitive-local). Mate pairing, sorting and duplicate removal were performed with Pairtools^57^, and contact matrices binned/visualised with Cooler^58^. A/B compartments were identified using the Eigenvector function from the cooltools package^59^. Insulation Valleys (IVs) were detected using HiCExplorer^30^ (hicFindTADs --thresholdComparisons 0.05 --delta 0.01 -- correctForMultipleTesting fdr -p 64 –minDepth 100000 --maxDepth 40000 --step 1500). Chromatin loops were called using Mustache^31^ (with the following parameters: -d 50000000 -r 5000 -pt 0.05 -st 0.7 -sz 1.0 -oc 5 -i 30 -p 32). Pileup analysis of loops was done with cooltools package^59^.

### Chip-seq and ATAC-seq processing

For the epigenetic analysis of *S.lycopersicum* and mESC, raw fastq files were mapped with Bowtie2^56^, sorted with SAMtools^60^, and deduplicated with “Picard Toolkit”, 2019, Broad Institute, GitHub Repository (https://broadinstitute.github.io/picard/) (with the parameters: MarkDuplicates -VALIDATION_STRINGENCY LENIENT -REMOVE_DUPLICATES true - ASSUME_SORTED true). Normalyzed coverage tracks were generated with deepTools2^61^ (with parameters: bamCoverage --numberOfProcessors 8 --binSize 10 --normalizeUsing BPM). Peaks were called with MACS2^62^. All datasets are listed in Supplementary Table 1.

### Transposable-element dating

TEs were annotated in each genome with RepeatMasker^63^ (using -species “viridiplantae”). Age was estimated from % divergence using a neutral substitution rate of 1.3 × 10⁻⁸ site⁻¹ yr⁻¹ for tomato:

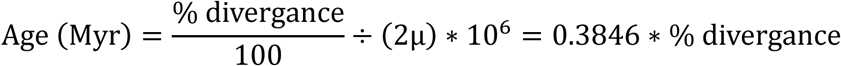

### Conservation of insulation valleys, loops and associated genes

For IV sequence conservation analysis, 5-kb windows centred on IV midpoints were aligned pairwise between species with Minimap2^54^ (minimap2 -ax asm20, ≥20 % identity). An IV was considered conserved only if its aligned window overlapped an IV in the target genome.

For chromatin loops sequence conservation analysis, both anchor windows (5 kb) of each loop were aligned as above; a loop was scored as conserved when the two aligned anchors formed a loop in the target genome. For aggregate-peak conservation, 5 kb windows centred on each *S. lycopersicum* loop anchor were lifted to the LA1589 and LA0716 assemblies with Minimap2^54^ (-ax asm20). These mapped coordinates were treated as candidate anchors in the wild relatives, and cooltools pileup was run on the three corresponding loop sets: the native *S.lycopersicum* loops and the LA1589 and LA0716 mapped counterparts, so that the resulting aggregate maps could be compared directly, revealing whether contact enrichment is preserved at orthologous positions across species.

For the analysis of Gene conservation, The M82 annotation was lifted over to LA1589 and LA0716 with Liftoff^64^. Genes whose TSS fell within ±2 kb of an IV or loop anchor were catalogued, and orthology was inferred from lift-over results. Functional enrichment was assessed with GOATOOLS^65^, using ITAG4.0 annotations^66^; shared enriched terms across species are reported.

### Mars-seq sequencing and analysis

Total RNA was extracted from young leaves of 4-week-old *S.lycopersicum* (cv. M82), *S.pimpinellifolium* (LA1589) and *S.pennellii* (LA0716) plants (ten biological replicates per species; each from a different plant) using the NucleoSpin RNA Plant kit (Macherey-Nagel). MARS-seq libraries were prepared as described by Keren-Shaul *et al*^44^ and sequenced on an Illumina NovaSeq X at the Nancy and Stephen Grand Israel National Center for Personalized Medicine (G-INCPM), Weizmann Institute of Science. Sequencing output was processed with UTAP^45^. For cross-species quantification, the *S.lycopersicum* M82 gene annotation (GFF3) was projected onto the de novo assemblies of *S.pimpinellifolium* LA1589 and *S.pennellii* LA0716 to generate species-specific references, which were then used to build the UTAP analysis templates.

