## Supplemental data for "Micro-C in *Solanum* Uncovers Conserved Genome Folding and Epigenetically Defined Loops with Bifunctional Enhancer-Silencer Activity"

Supplementary figures:

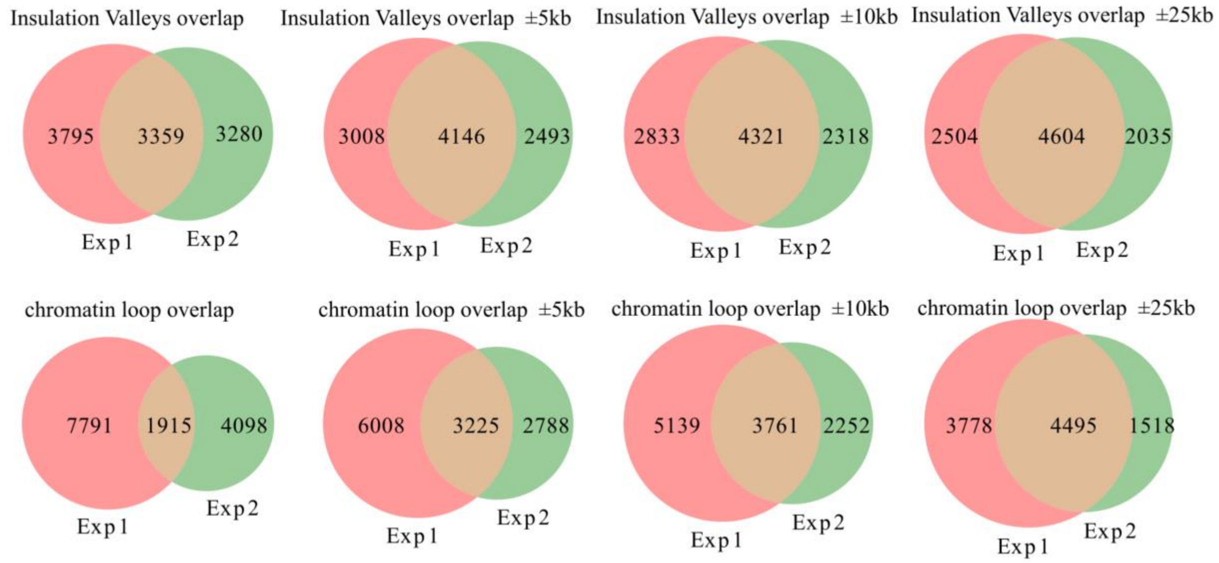


Supplementary figure 1: **Overlap of insulation valleys and chromatin loops between two experimental replicates of Solanum lycopersicum Micro-C.** Top panel: Insulation-valley overlap at increasing tolerance windows: exact, ±5 kb, ±10 kb and ±25 kb (from left to right). Bottom panel: Chromatin-loop overlap between the same replicates, using the same series of tolerance windows- exact, ±5 kb, ±10 kb and ±25 kb (from left to right).

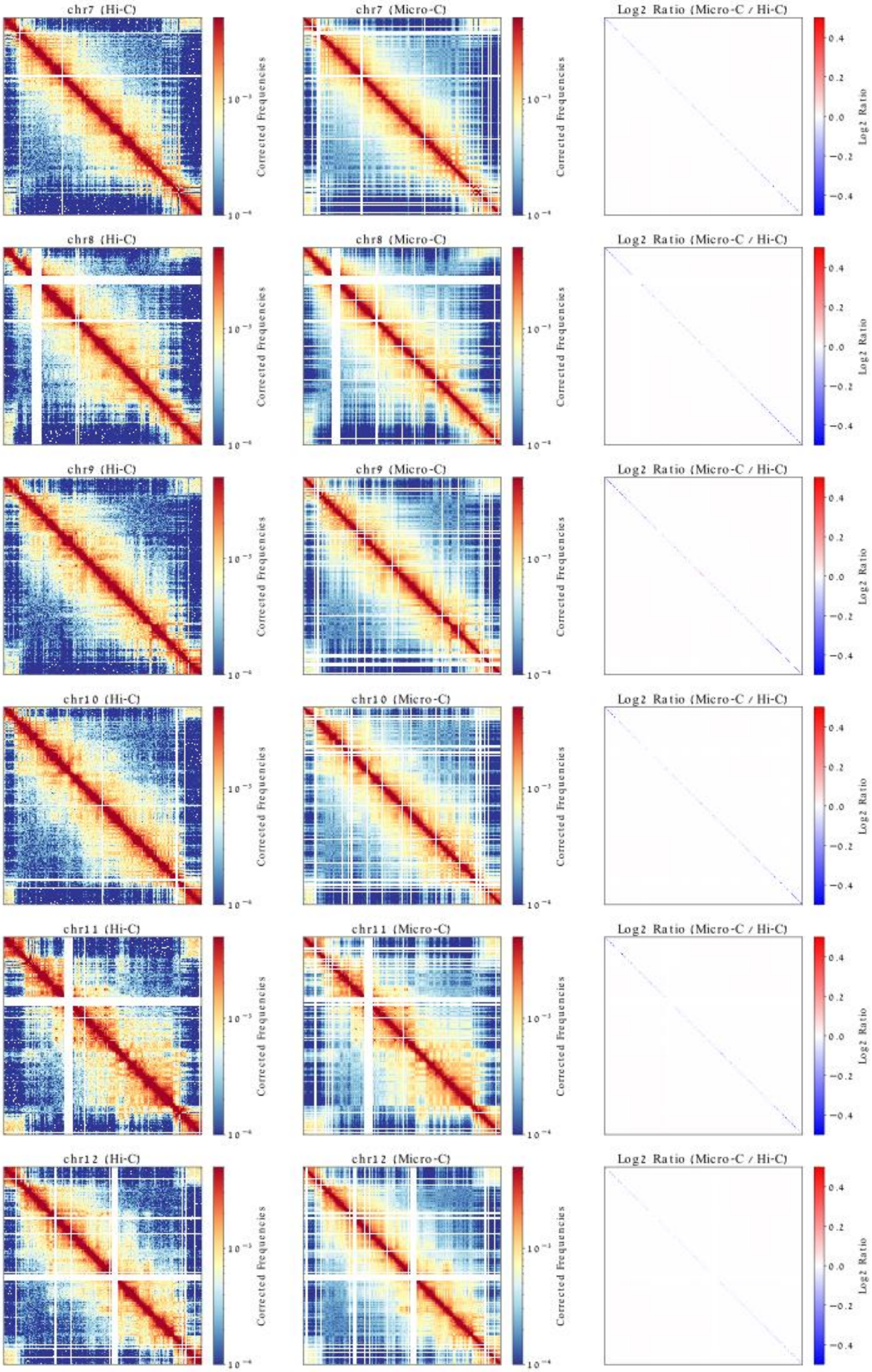
chr7 (Hi-C)
Corrected Frequencies
chr7 (Micro-C)
Corrected Frequencies
Log2 Ratio (Micro-C / Hi-C)
Log2 Ratio
chr8 (Hi-C)
Corrected Frequencies
chr8 (Micro-C)
Corrected Frequencies
Log2 Ratio (Micro-C / Hi-C)
Log2 Ratio
chr9 (Hi-C)
Corrected Frequencies
chr9 (Micro-C)
Corrected Frequencies
Log2 Ratio (Micro-C / Hi-C)
Log2 Ratio
chr10 (Hi-C)
Corrected Frequencies
chr10 (Micro-C)
Corrected Frequencies
Log2 Ratio (Micro-C / Hi-C)
Log2 Ratio
chr11 (Hi-C)
Corrected Frequencies
chr11 (Micro-C)
Corrected Frequencies
Log2 Ratio (Micro-C / Hi-C)
Log2 Ratio
chr12 (Hi-C)
Corrected Frequencies
chr12 (Micro-C)
Corrected Frequencies
Log2 Ratio (Micro-C / Hi-C)
Log2 Ratio

Similarity Metrics for Whole Chromosomes

| Chromosome | Pearson Corr | Spearman Corr | MSE | SSIM |
|---|---|---|---|---|
| chr1 | 0.919 | 0.858 | 0.0 | 0.996 |
| chr2 | 0.874 | 0.911 | 0.0 | 0.992 |
| chr3 | 0.919 | 0.902 | 0.0 | 0.993 |
| chr4 | 0.919 | 0.889 | 0.0 | 0.994 |
| chr5 | 0.934 | 0.879 | 0.0 | 0.995 |
| chr6 | 0.913 | 0.923 | 0.0 | 0.993 |
| chr7 | 0.921 | 0.892 | 0.0 | 0.994 |
| chr8 | 0.926 | 0.891 | 0.0 | 0.996 |
| chr9 | 0.923 | 0.873 | 0.0 | 0.994 |
| chr10 | 0.932 | 0.887 | 0.0 | 0.994 |
| chr11 | 0.942 | 0.898 | 0.0 | 0.996 |
| chr12 | 0.932 | 0.882 | 0.0 | 0.996 |

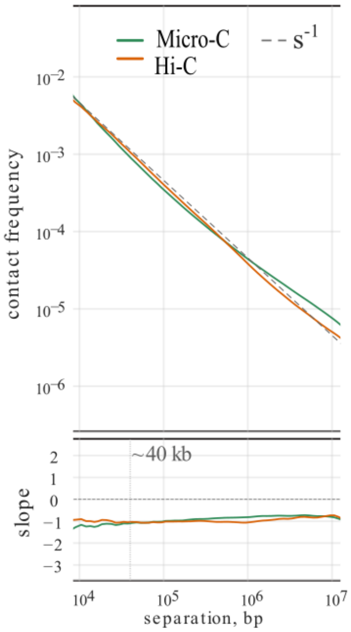


Supplementary figure 2: **Comparison of Hi-C and Micro-C heatmaps across all chromosomes in *S.lycopersicum.*** Each panel corresponds to one chromosome: Hi-C data from Huang Y. *et al*[16] (left), Micro-C (center), and $\log_2$ ratio (Micro-C/Hi-C) (right). a. The accompanying table summarizes Pearson's and Spearman's correlation coefficients, Mean Squared Error (MSE), and Structural Similarity Index (SSIM) for each chromosome at the whole-chromosome level. B. Interaction probability by distance (top) and its first derivative (bottom) for M82 measured by Micro-C and Hi-C. Both assays follow an $\sim s^{-1}$ decay and share a trough near ~40 kb, indicating similar meso-scale folding. Micro-C reports higher contact frequency at nucleosome scales (~1–8 kb) and retains stronger, smoother signal at long distances (>1 Mb), giving a shallower minimum and slopes closer to −1; Hi-C shows a steeper short-range drop and a faster decay in the tail.

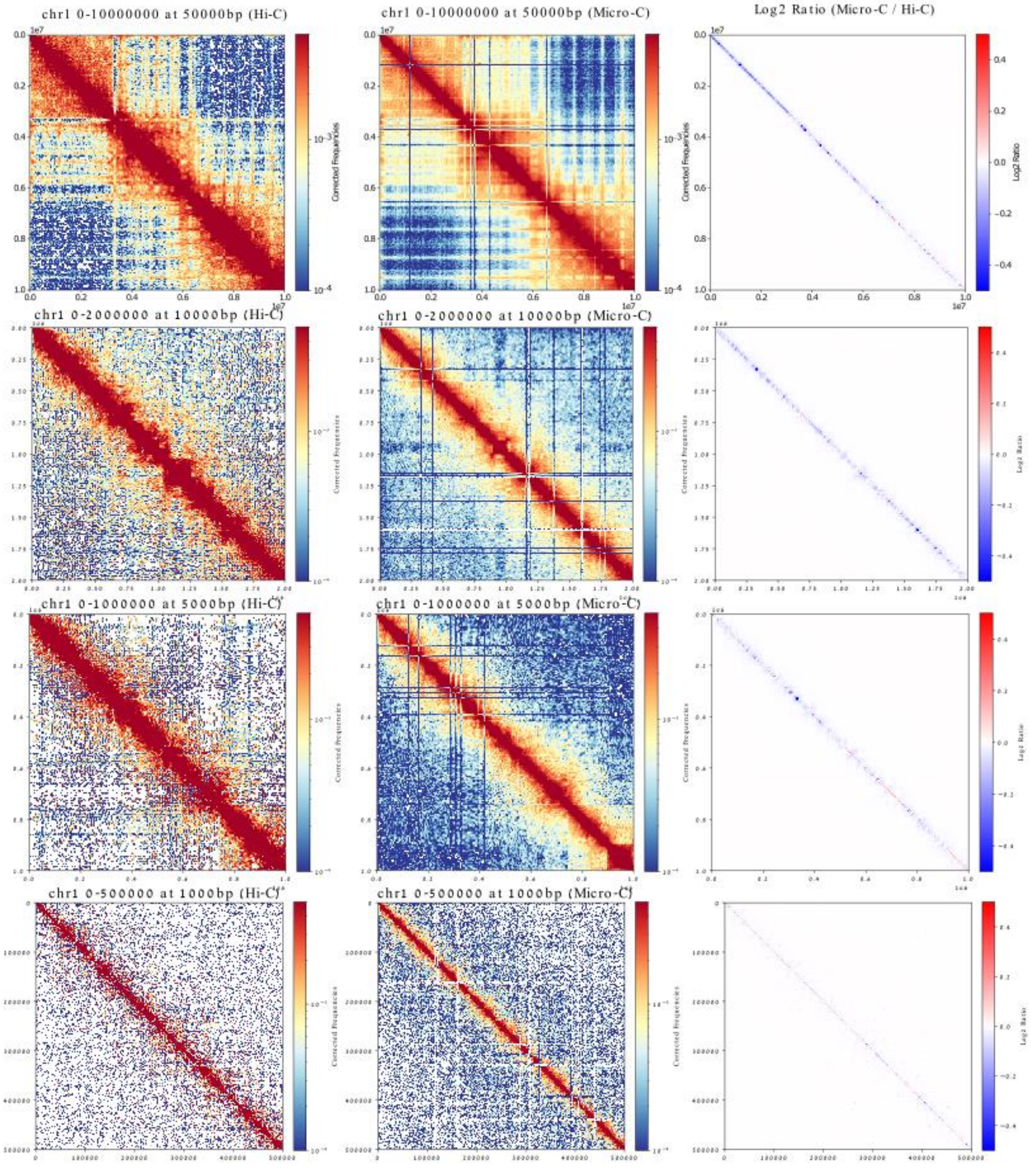


Similarity Metrics for Chromosome 1 at Multiple Resolutions

| Region | Pearson Corr | Spearman Corr | MSE | SSIM |
|---|---|---|---|---|
| chr1:0-10000000 (50000bp) | 0.927 | 0.903 | 0.0 | 0.976 |
| chr1:0-2000000 (10000bp) | 0.882 | 0.563 | 0.0 | 0.962 |
| chr1:0-1000000 (5000bp) | 0.816 | 0.564 | 0.0 | 0.936 |
| chr1:0-500000 (1000bp) | 0.672 | 0.337 | 0.0 | 0.895 |

Supplementary figure 3: **Comparison of Hi-C and Micro-C heatmaps at multiple resolutions in *S.lycopersicum*.** Each panel shows a segment of chromosome 1 at a specified bin size, with Hi-C data from Huang et al.[16] (left), Micro-C (center), and the $\log_2$ ratio (Micro-C/Hi-C) (right). The accompanying table summarizes Pearson's and Spearman's correlation coefficients, mean squared error (MSE) and structural similarity index (SSIM) for each resolution.

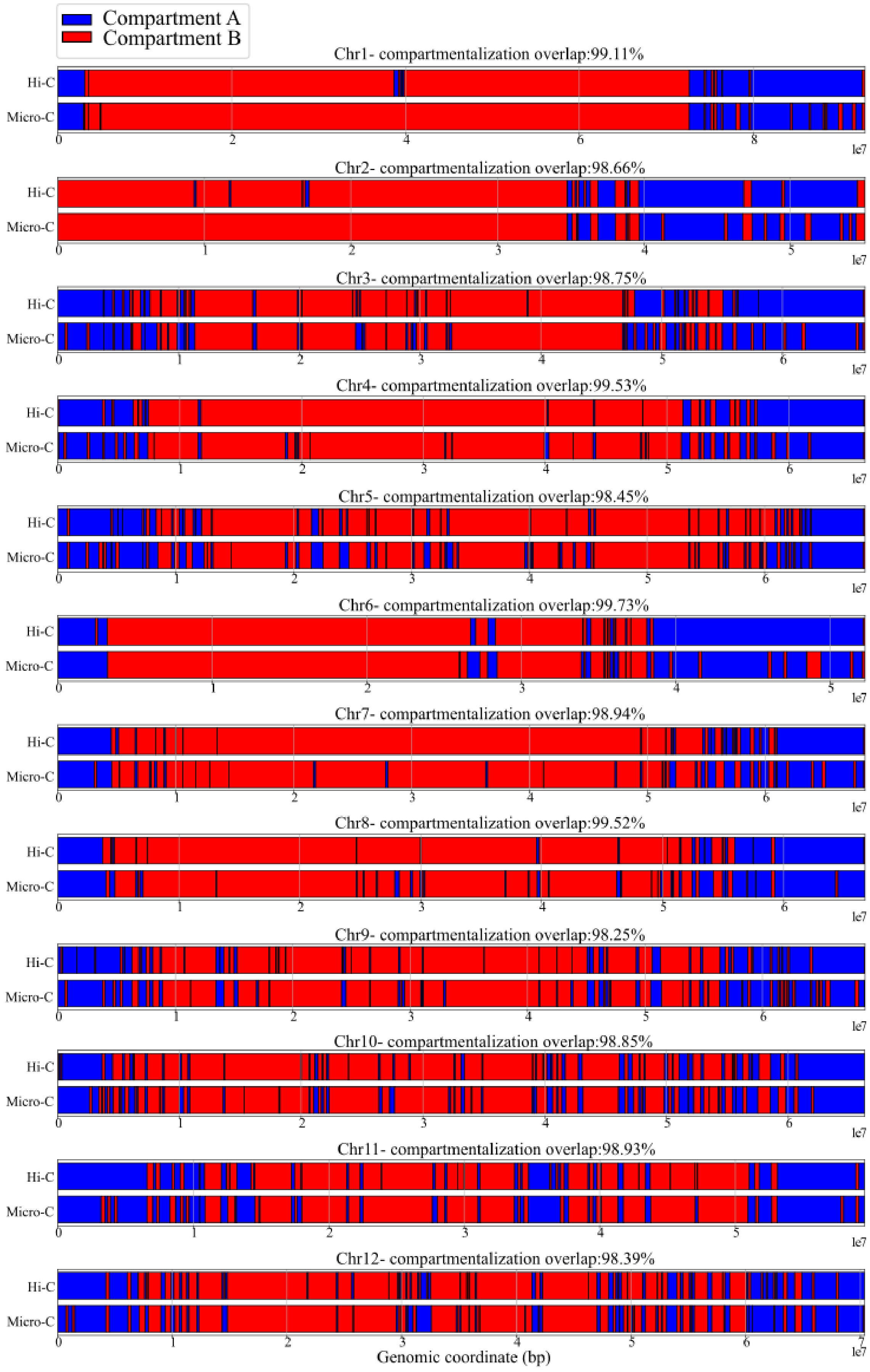
Compartment A
Compartment B
Chr1- compartmentalization overlap:99.11%
Chr2- compartmentalization overlap:98.66%
Chr3- compartmentalization overlap:98.75%
Chr4- compartmentalization overlap:99.53%
Chr5- compartmentalization overlap:98.45%
Chr6- compartmentalization overlap:99.73%
Chr7- compartmentalization overlap:98.94%
Chr8- compartmentalization overlap:99.52%
Chr9- compartmentalization overlap:98.25%
Chr10- compartmentalization overlap:98.85%
Chr11- compartmentalization overlap:98.93%
Chr12- compartmentalization overlap:98.39%
Hi-C
Micro-C
1e7
Genomic coordinate (bp)

Supplementary Figure 4: **Chromosome-wide compartment annotation from Hi-C versus Micro-C in *S.lycopersicum*.** Compartment A/B assignments were derived by eigenvector decomposition of each contact heatmap. For each chromosome, the top track shows Hi-C compartments and the bottom track shows Micro-C compartments (A in blue, B in red). The percentage overlap between Hi-C and Micro-C compartment calls is indicated above each chromosome.

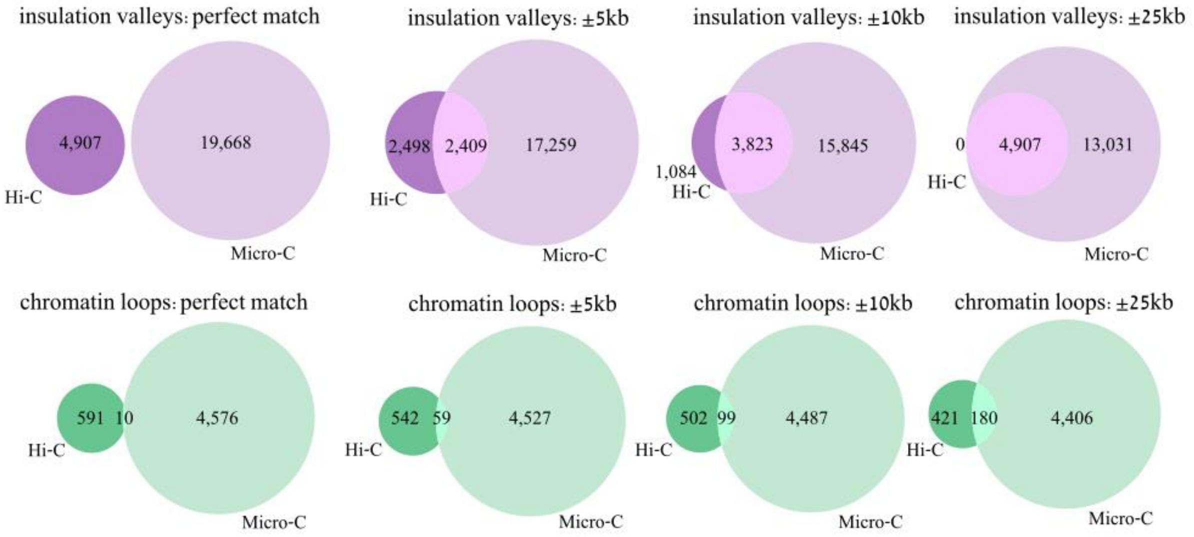


Supplementary Figure 5: **Overlap of insulation valleys and chromatin loops between Hi-C and Micro-C in *S.lycopersicum*.** Top panel: Insulation-valley overlap evaluated at increasing tolerance windows - exact match (0 bp), ±5 kb, ±10 kb and ±25 kb.

Bottom panel: Chromatin-loop overlap evaluated at the same series of tolerance windows - exact match (0 bp), ±5 kb, ±10 kb and ±25 kb.

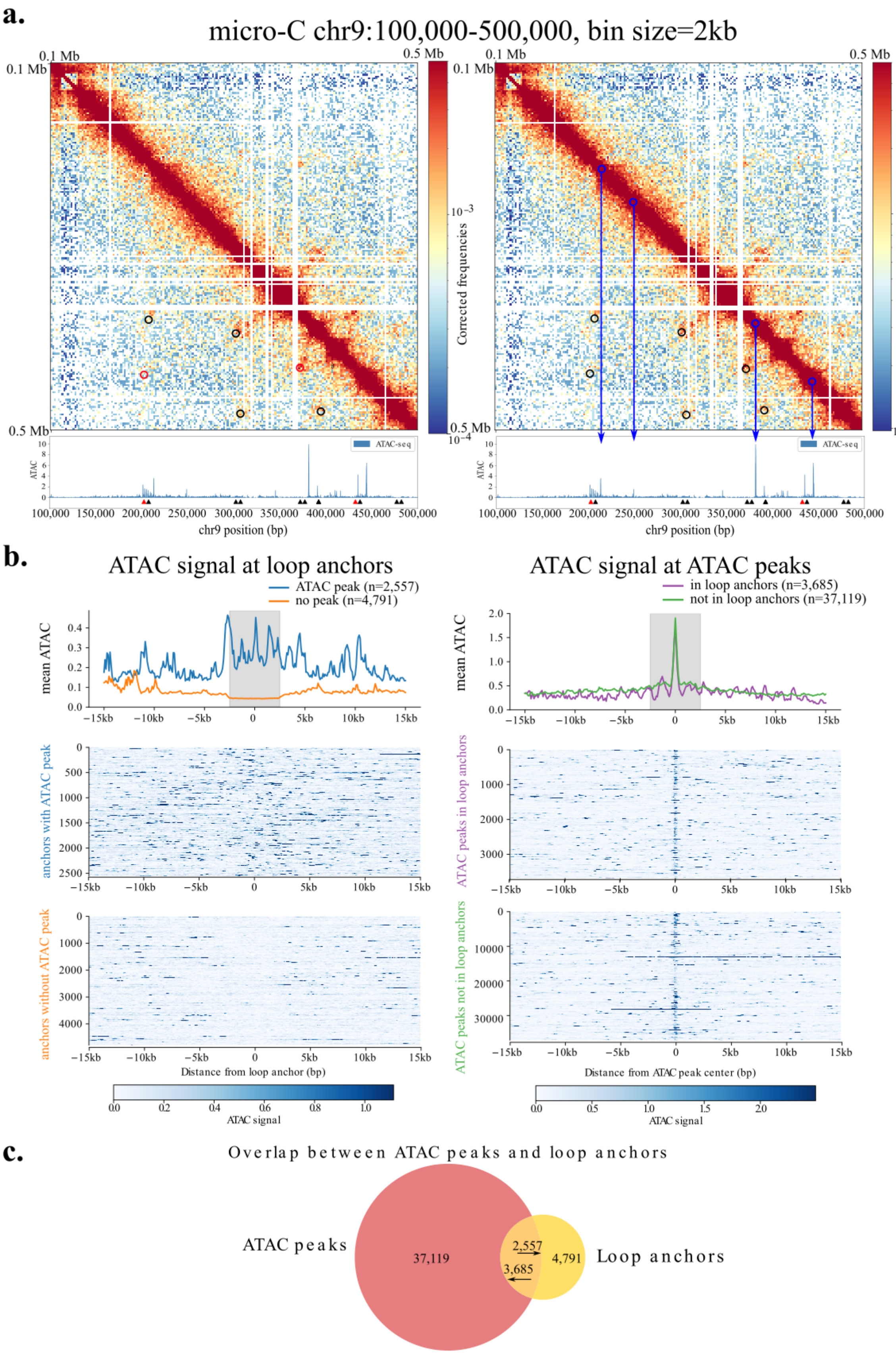
a.
micro-C chr9:100,000-500,000, bin size=2kb
0.1 Mb
0.5 Mb
Corrected frequencies
$10^{-3}$
$10^{-4}$
ATAC-seq
ATAC
chr9 position (bp)
100,000 150,000 200,000 250,000 300,000 350,000 400,000 450,000 500,000
b.
ATAC signal at loop anchors
ATAC peak (n=2,557)
no peak (n=4,791)
mean ATAC
anchors with ATAC peak
anchors without ATAC peak
Distance from loop anchor (bp)
ATAC signal
ATAC signal at ATAC peaks
in loop anchors (n=3,685)
not in loop anchors (n=37,119)
ATAC peaks in loop anchors
ATAC peaks not in loop anchors
Distance from ATAC peak center (bp)
−15kb −10kb −5kb 0 5kb 10kb 15kb
c.
Overlap between ATAC peaks and loop anchors
ATAC peaks
37,119
2,557
3,685
4,791
Loop anchors

Supplementary Figure 6: **Micro-C maps reveal chromatin loop anchors without ATAC-seq signal and ATAC peaks not involved in loop formation.** a. Micro-C heatmap of *chr9:100,000–500,000* at 2 kb resolution. Right panel: Loop anchors overlapping ATAC-seq peaks are marked in red; anchors without ATAC peaks are marked in black. The corresponding ATAC peaks are indicated by red and black arrows, respectively. Left panel: All detected loops are shown in black. ATAC peaks not overlapping loop anchors are highlighted with blue circles, with arrows pointing to the corresponding peaks. b. Right: average ATAC-seq signal at loop anchors with (blue) and without (orange) overlapping ATAC peaks. Left: average ATAC-seq signal at peaks that overlap (purple) or do not overlap (green) chromatin loop anchors. C. overlap between ATAC peaks and loop anchors. 2,557 of loop anchors containing ATAC peak and 3,685 of ATAC peaks are located at loop anchors.

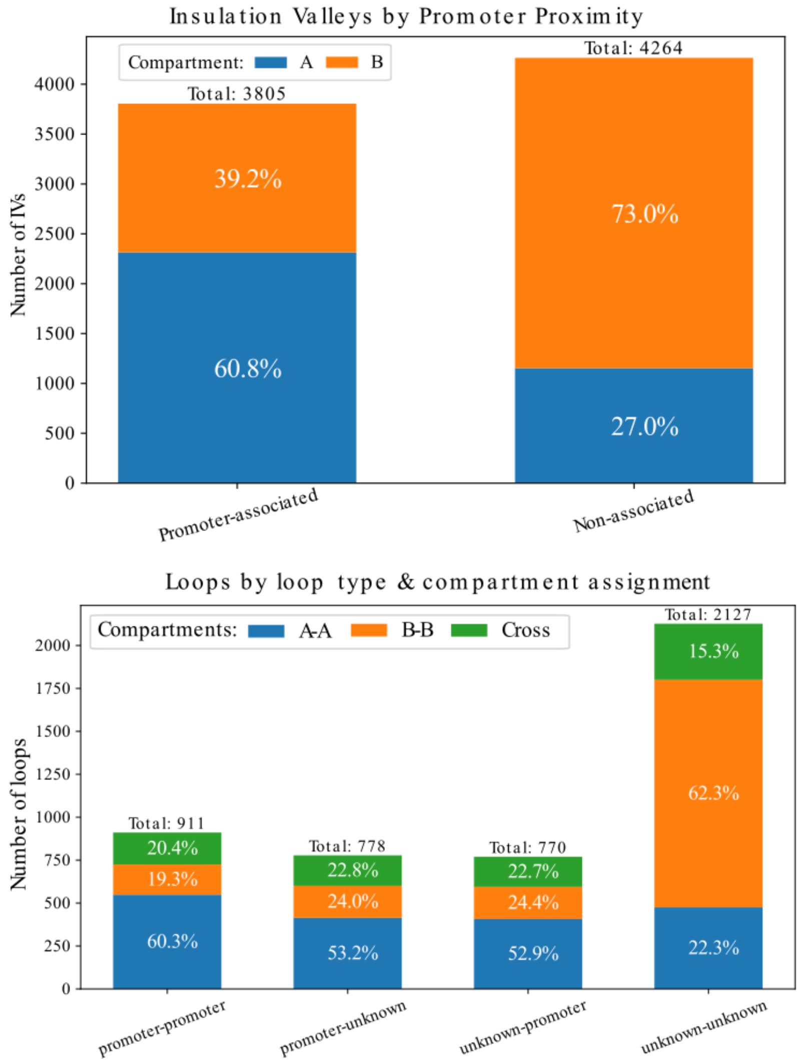


Supplementary Figure 7: **Insulation valleys and chromatin loop classification in *S.lycopersicum*.**

Top: Insulation valleys were split into promoter-associated (IV centers within ±2 kb of a transcription start site) and non-promoter IVs; each group is further subdivided by chromatin compartment. Bottom: Chromatin loops were classified by promoter proximity at their anchors- anchors whose centers lie within ±2 kb of a TSS are “promoter anchors”-resulting in promoter-promoter (P-P), promoter–unknown (P-U), unknown-promoter (U-P), and unknown-unknown (U-U) loops; each loop class is likewise subdivided by chromatin compartment.

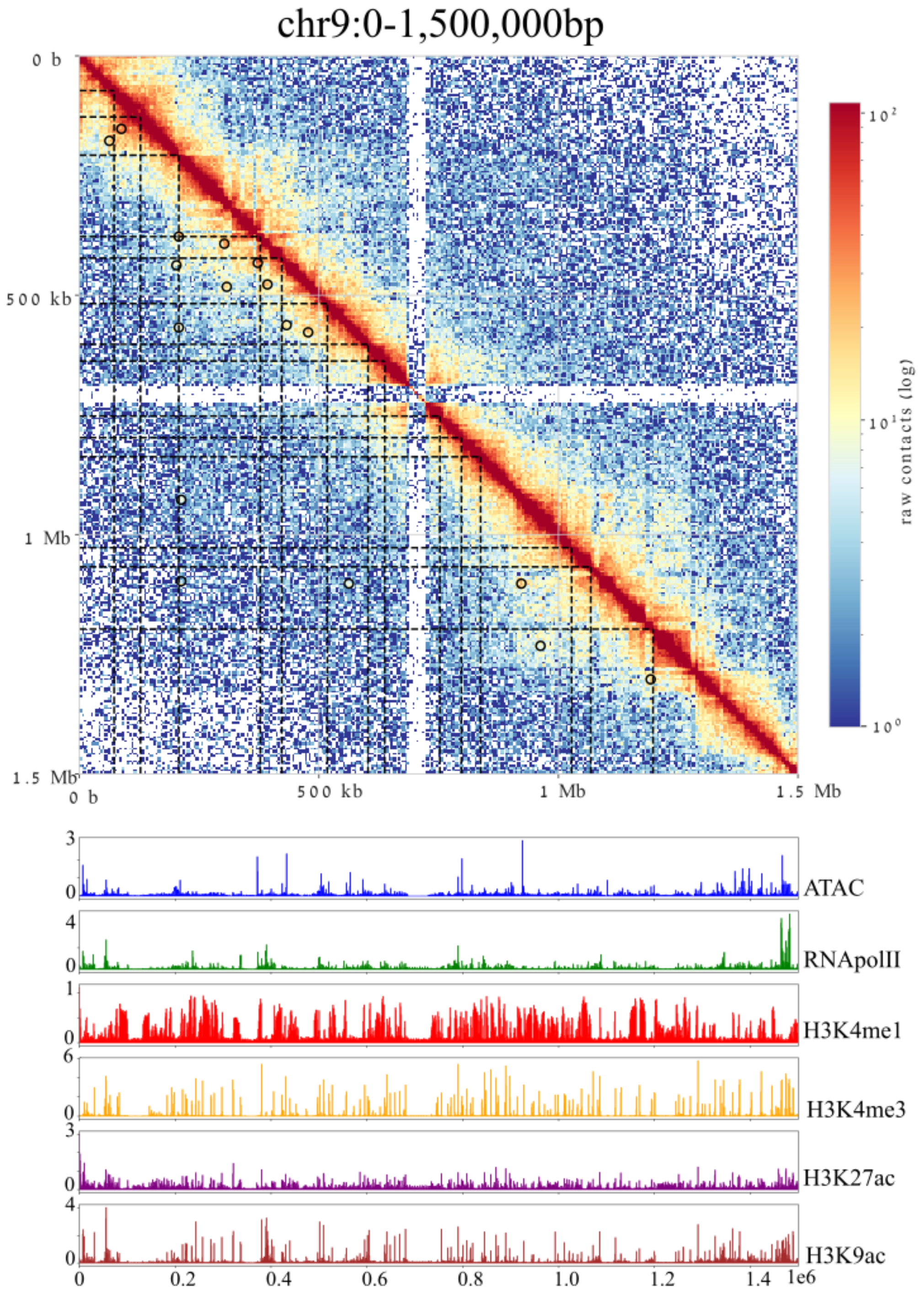


Supplementary Figure 8: **Micro-C contact map and ChIP-seq profiles on *S.lycopersicum* chromosome 9:0-1,500,00bp.** Top panel: Micro-C heatmap (0–1.5 Mb) with insulation valleys marked by black dashed lines and loop anchors highlighted with black circles. Bottom panel: Signal tracks for ATAC-seq, RNA polymerase II, H3K4me1, H3K4me3, H3K27ac and H3K9ac across the same genomic interval.

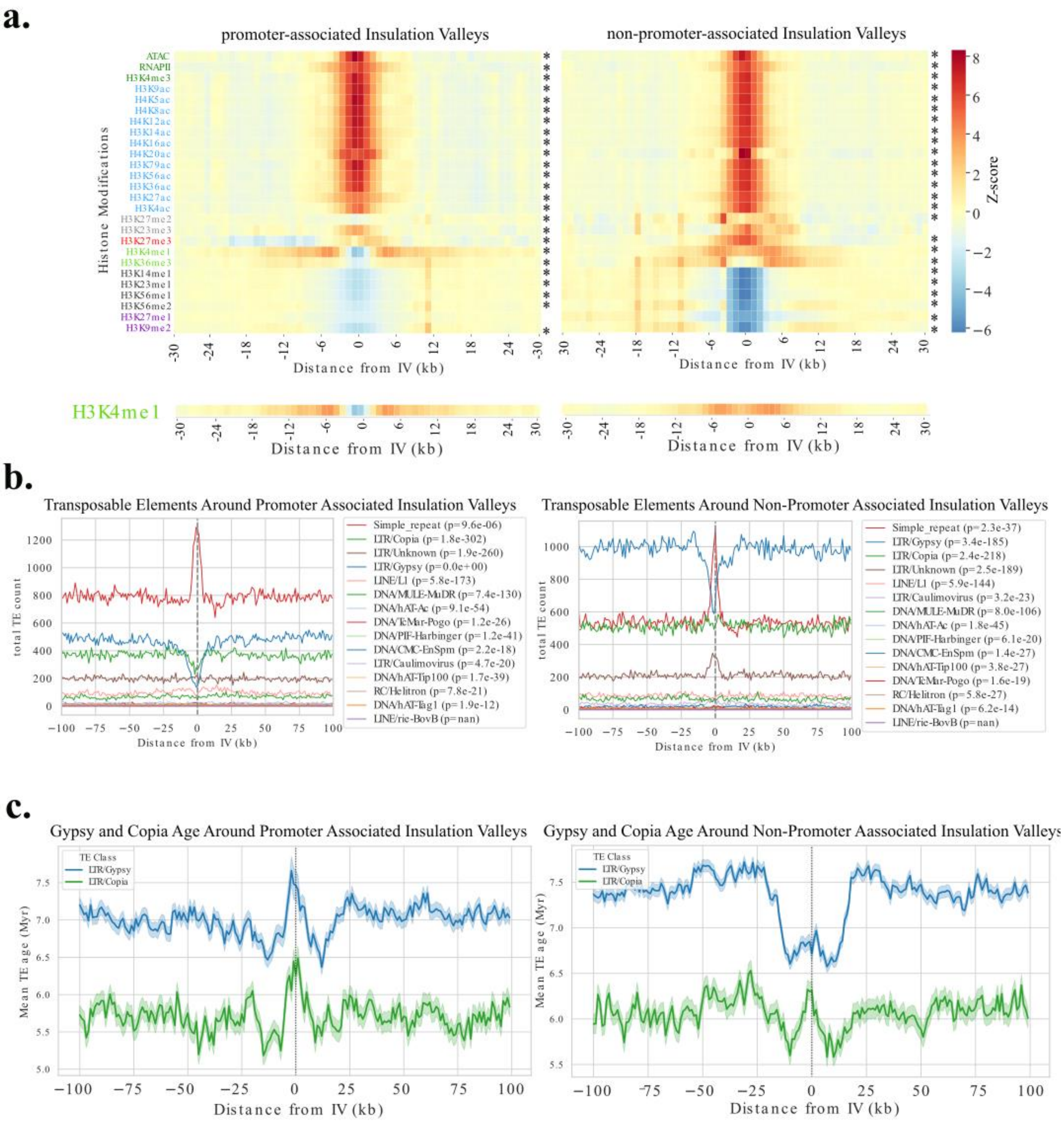


Supplementary Figure 9 **Histone modifications and transposable-element landscapes at insulation valleys.** a Average histone-modification profiles around promoter-associated versus non-promoter IVs (±100 kb from IV centers). Top: all tested histone marks; asterisks to the left of each panel indicate significant enrichment or depletion at the IV center (±2.5 kb) compared to distal flanks (±15-30 kb), based on a paired Wilcoxon test ($p < 0.01$). Bottom: H3K4me1 alone, shown separately for clarity. b. Genome-wide density profiles of major transposable-element (TE) classes in ±100 kb windows centered on promoter-associated and non-promoter IVs. P-values indicate the significance of TE depletion or enrichment at the IV center (±2.5 kb) relative to distal flanks (±80–100 kb), using a paired Wilcoxon test. c. Mean ages of Gypsy and Copia elements in ±100 kb around IV centers; shaded regions represent the standard error of the mean (SEM).

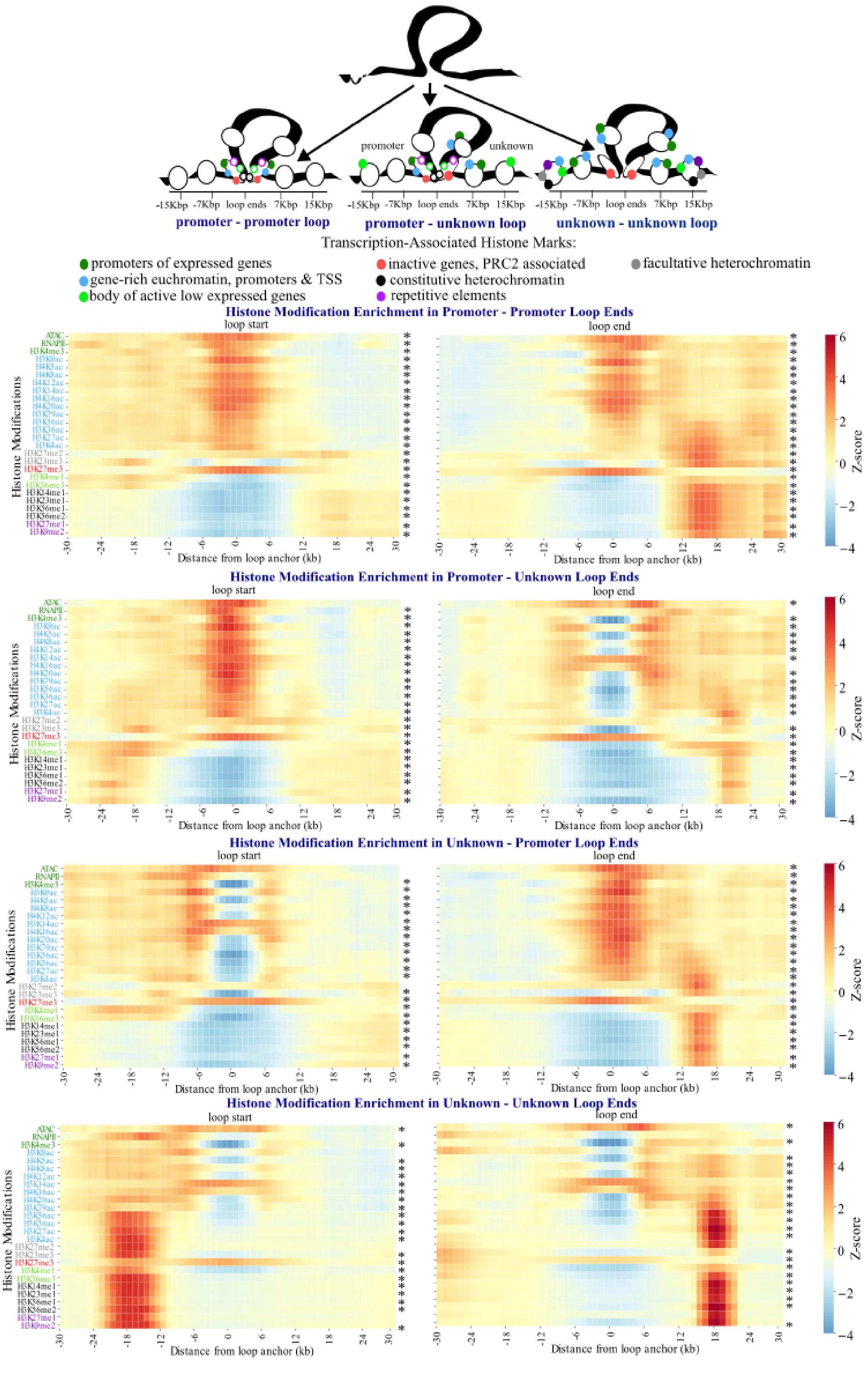

promoter
unknown
-15Kbp -7Kbp loop ends 7Kbp 15Kbp
promoter - promoter loop
promoter - unknown loop
unknown - unknown loop
Transcription-Associated Histone Marks:
promoters of expressed genes
gene-rich euchromatin, promoters & TSS
body of active low expressed genes
inactive genes, PRC2 associated
constitutive heterochromatin
repetitive elements
facultative heterochromatin
Histone Modification Enrichment in Promoter - Promoter Loop Ends
Histone Modification Enrichment in Promoter - Unknown Loop Ends
Histone Modification Enrichment in Unknown - Promoter Loop Ends
Histone Modification Enrichment in Unknown - Unknown Loop Ends
loop start
loop end
Histone Modifications
Distance from loop anchor (kb)
Z-score

Supplementary Figure 10: **Histone modification patterns at chromatin loop anchors in *S.lycopersicum*. Top panel:** Schematic models of histone modification landscapes at promoter-promoter (P-P), promoter-unknown (P-U; analogous to U-P) and unknown-unknown (U-U) loop anchors. **Bottom panel:** Z-score-normalized ChIP-seq signal (range -4 to +6) for each loop category. Histone marks are grouped and color-coded by functional annotation: **Promoter-associated (dark green):** ATAC-seq, RNA polymerase II, H3K4me3, **Transcription start site-associated (light blue):** H3K9ac, H4K5ac, H4K8ac, H4K12ac, H3K14ac, H4K16ac, H4K20ac, H3K79ac, H3K56ac, H3K36ac, H3K27ac, H3K4ac, **Gene-body/low-expression (light green):** H3K4me1, H3K36me3, **Facultative heterochromatin (grey):** H3K27me2, H3K23me3, **PRC2-associated (red):** H3K27me3, **Constitutive heterochromatin (black):** H3K14me1, H3K23me1, H3K56me1, H3K56me2, **Repeat-associated (purple):** H3K27me1, H3K9me2. asterisks to the left of each panel indicate significant enrichment or depletion at the loop anchor center (±2.5 kb) compared to distal flanks (±15-30 kb), based on a paired Wilcoxon test ($p < 0.01$).

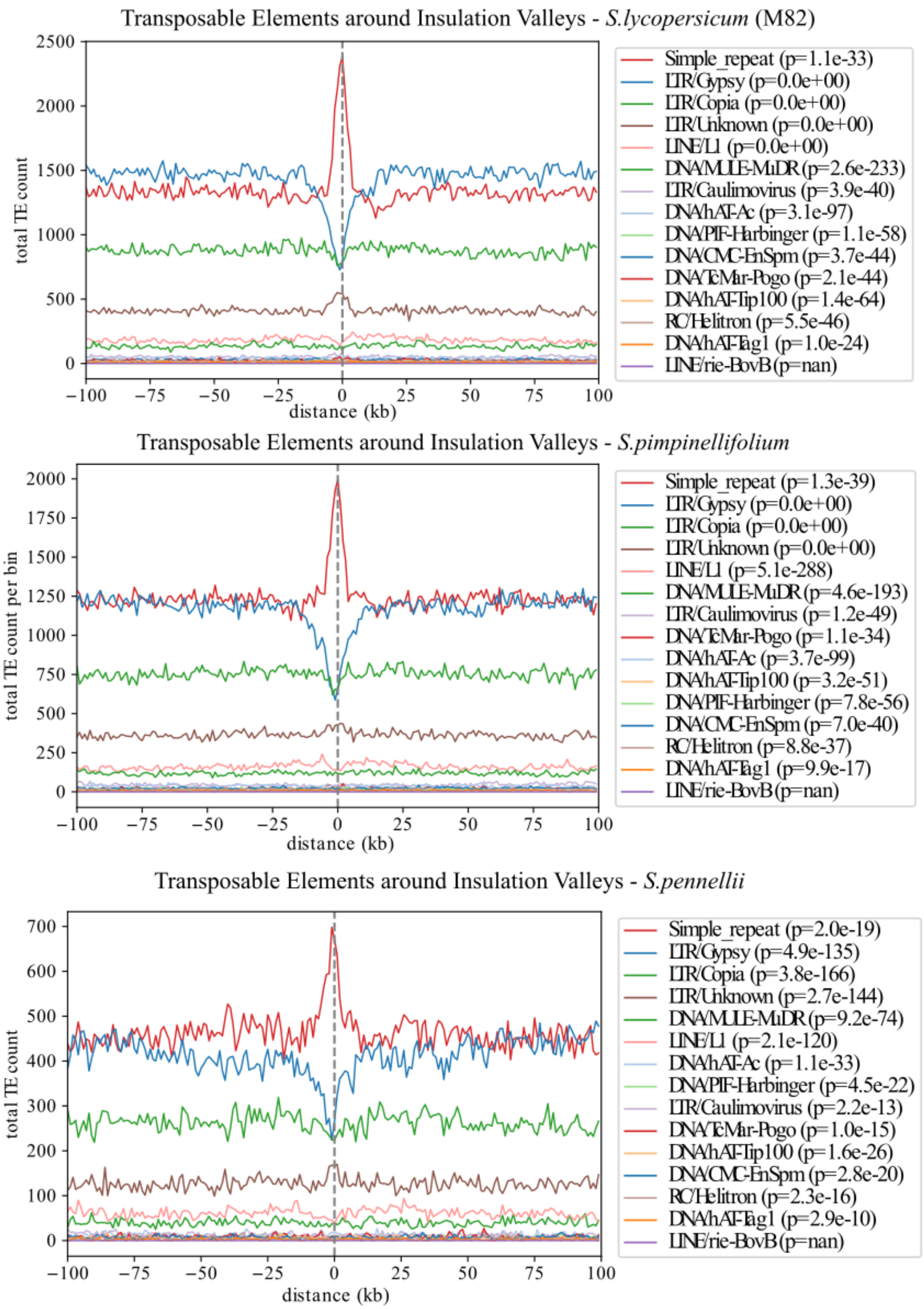


Supplementary Figure 11: **Transposable-element distribution around insulation valleys in three *Solanum* species.** Top panel: *S.lycopersicum*, Middle panel: *S.pimpinellifolium*, Bottom panel: *S.pennellii.* Each profile shows the genome-wide density of transposable elements relative to insulation-valley centers (dashed black line at 0). Wilcoxon signed-rank test p-values (in parentheses) indicate the significance of enrichment (positive deviation) or depletion (negative deviation) of each element class at valley centers compared to the flanking regions.

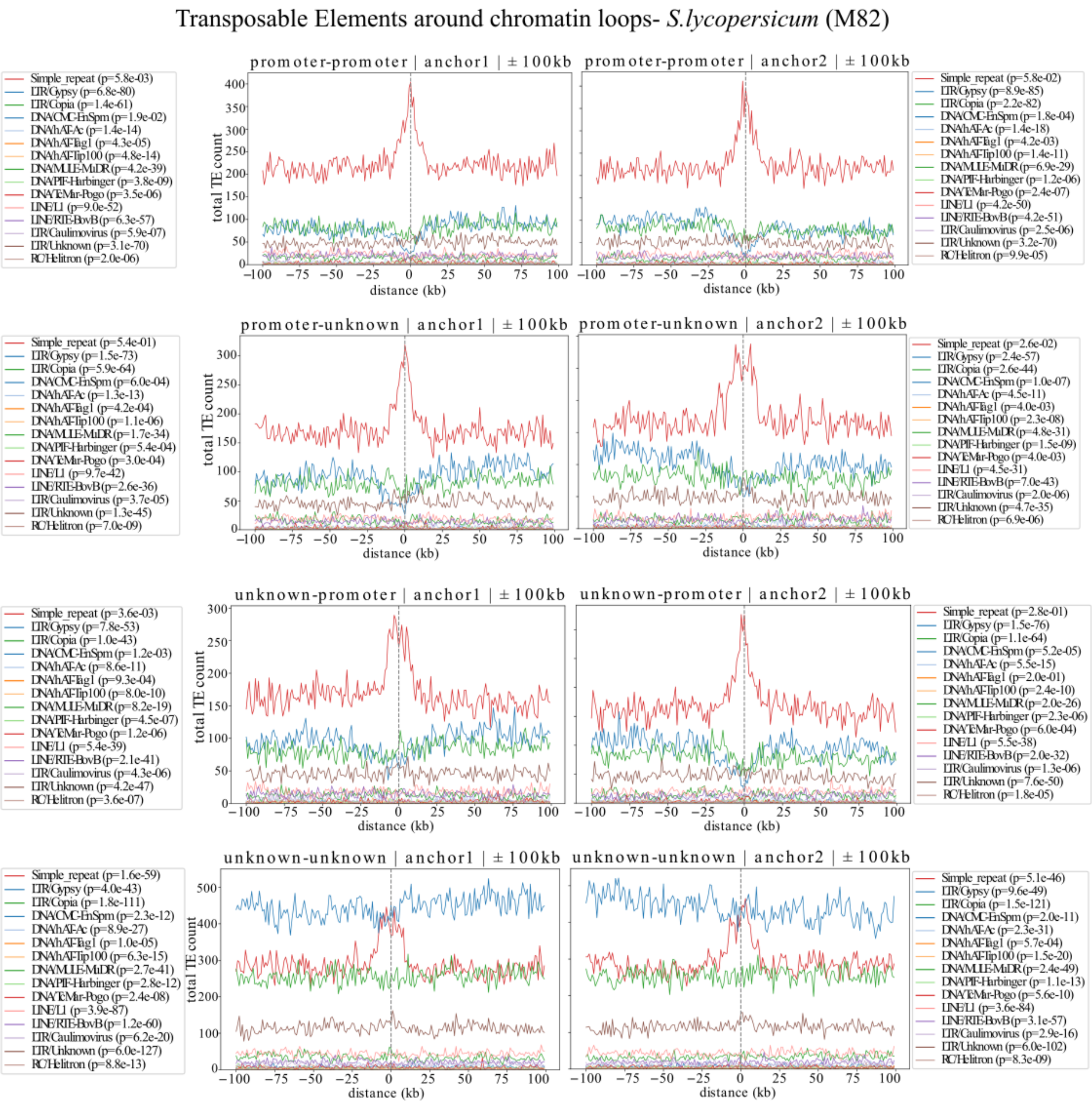


Supplementary Figure 12: **Transposable-element distribution around chromatin loop anchors in *S.lycopersicum*.** Panels show (from top to bottom): promoter-promoter loops, promoter-unknown loops, unknown-promoter loops, and unknown-unknown loops. Each profile depicts the density of transposable elements in a ±100 kb window centered on loop anchors (dashed black line at 0). Wilcoxon signed-rank test p-values (in parentheses) indicate the significance of enrichment (positive deviation) or depletion (negative deviation) of each element class at the anchor center compared to the flanking regions.

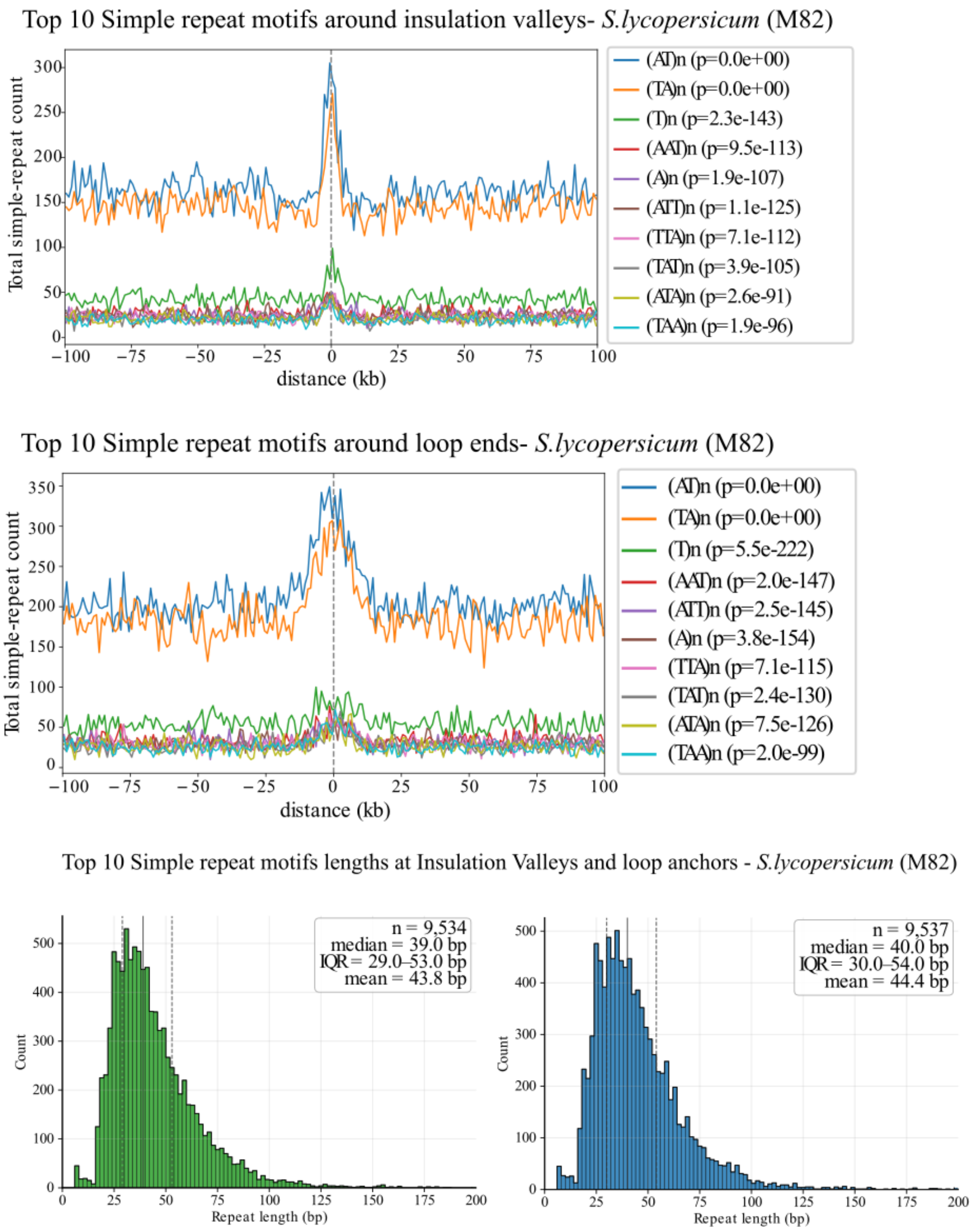


Supplementary Figure 13: **Simple repeats distribution around insulation valleys and chromatin loop anchors in *S.lycopersicum*.** Top panel: distribution around insulation valley centers. Middle panel: distribution around loop anchors. Each profile depicts the density of simple repeats in a ±100 kb window centered on either insulation valleys or loop anchors (dashed black line at 0). Wilcoxon signed-rank test p-values (in parentheses) indicate the significance of enrichment (positive deviation) or depletion (negative deviation) of simple repeats at the center compared to the flanking regions. Length distributions of simple repeats at IVs (left, green) and loop anchors (right, blue) limited to ≤200 bp; boxes report sample size (*n*), median, mean, and interquartile range (IQR = Q1–Q3).

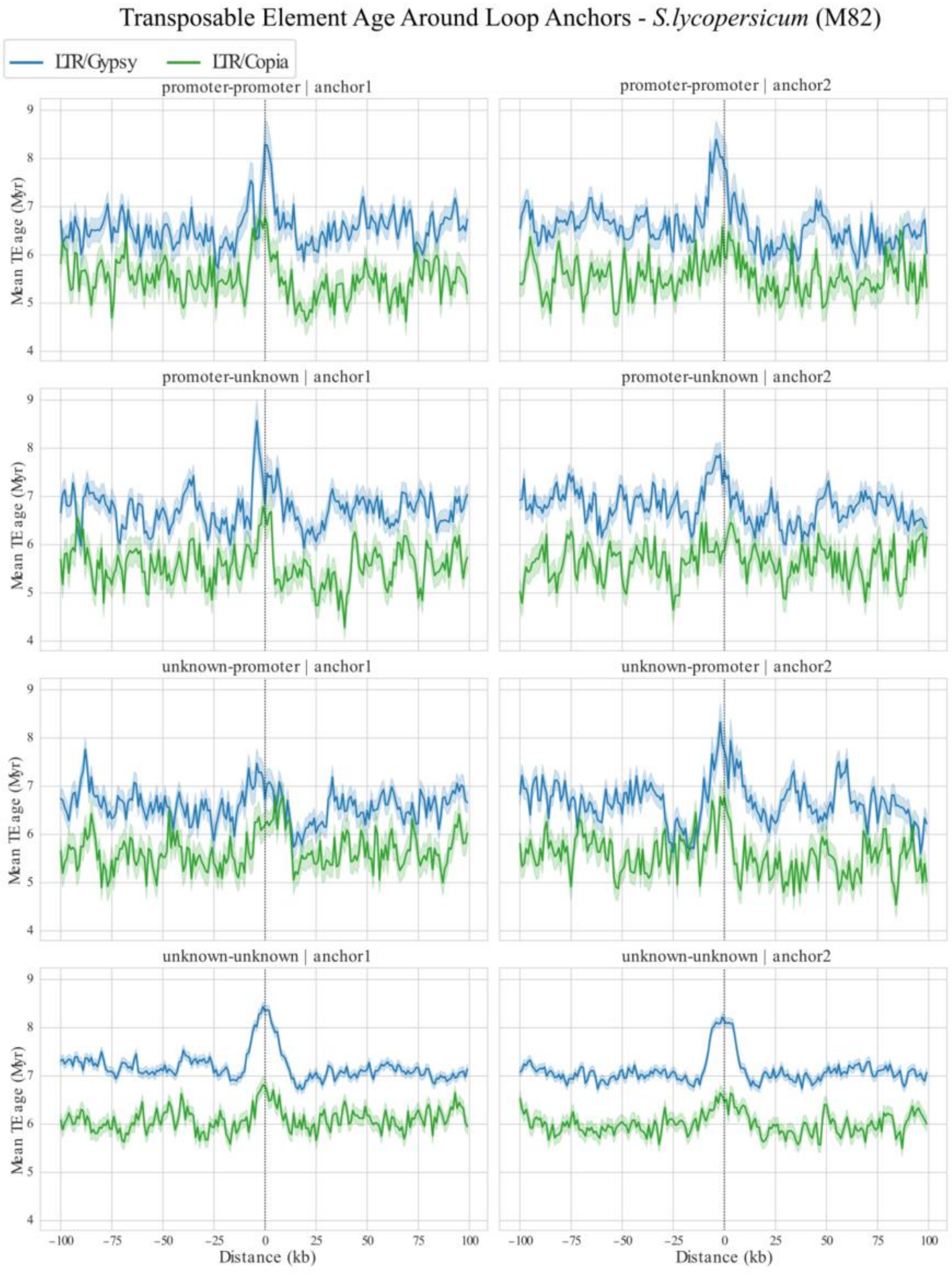


Supplementary Figure 14: **Mean age distribution of Gypsy and Copia elements around chromatin loop anchors in *S.lycopersicum*.** Panels (top to bottom) correspond to promoter-promoter loops, promoter-unknown loops, unknown-promoter loops, and unknown-unknown loops. Each profile shows the mean age of Gypsy and Copia transposable elements in a ±100 kb window centered on loop anchors (dashed black line at 0), with shaded regions representing the standard error of the mean (SEM).

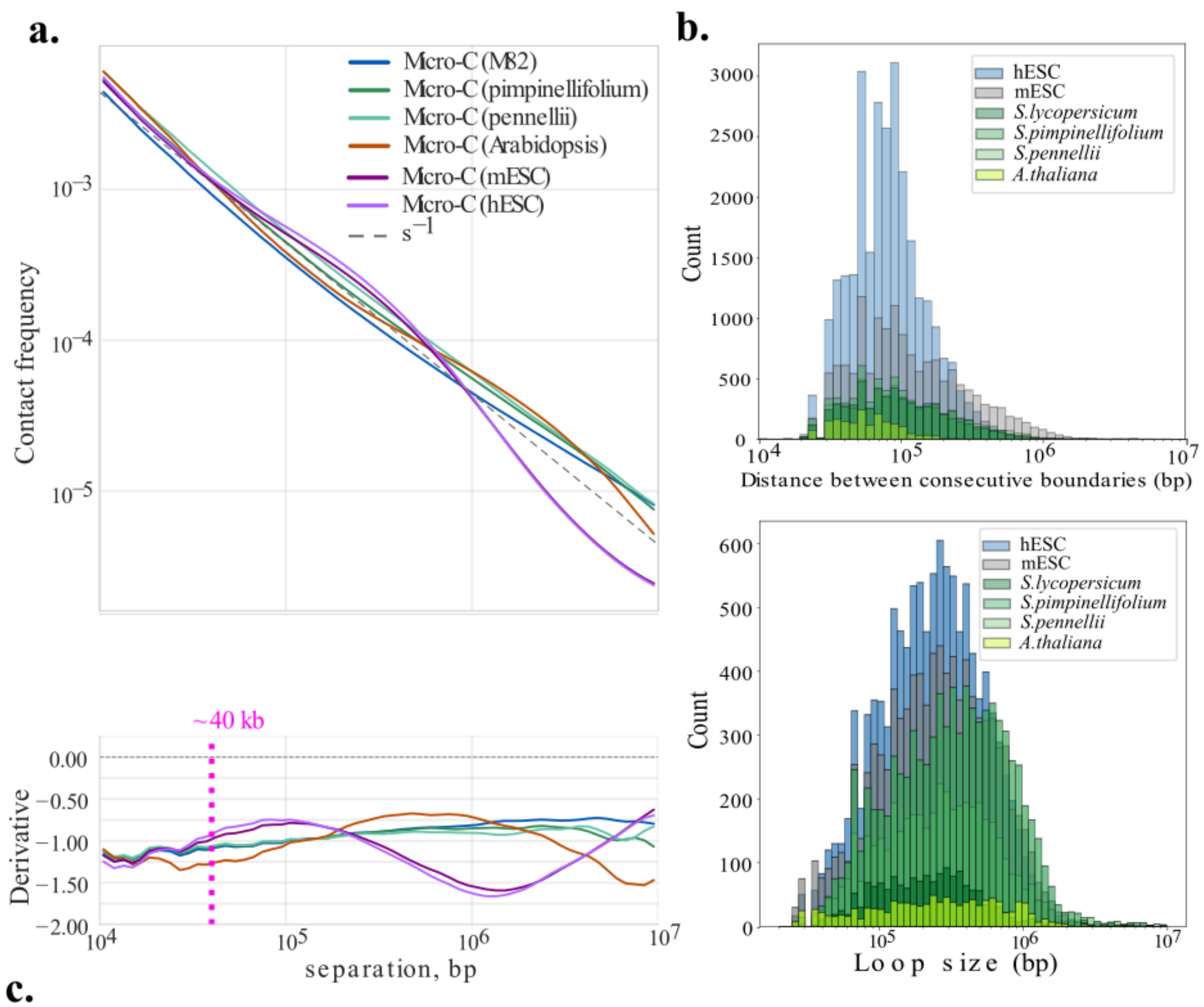


c.

| | *S.lycopersicum* (M82) | *S.pimpinellifolium* | *S.pennellii* | *A.thaliana* | mESC | hESC |
|---|---|---|---|---|---|---|
| Genome size (bp)* | 801,960,645 | 1,055,587,603 | 1,154,407,874 | 119,668,634 | 2,755,513,859 | 3,088,269,832 |
| Number of IVs** | 5905 | 6450 | 6023 | 1785 | 14,585 | 28,254 |
| Average distance between IVs** | 135,349 | 125,909 | 153,458 | 66,193 | 176,246 | 104,695 |
| N50 of distance between IVs** | 195,000 | 180,000 | 240,000 | 75,000 | 300,000 | 120,000 |
| Number of loops** | 1,643 | 8,385 | 4,658 | 1,138 | 9,546 | 11,570 |
| Average loop size** | 301,397 | 461,147 | 404,684 | 403,884 | 356,214 | 318,725 |
| N50 of loop size** | 410,000 | 710,000 | 615,000 | 255,000 | 545,000 | 455,000 |

* Genome size refers only to the combined length of chromosomes/contigs.
**Normalized by coverage.

Supplementary Figure 15: **Size distributions of insulation valleys and chromatin loops in plant and mammalian cells.** a. Depth-normalized 1-kb Micro-C interaction probability *P(s)* versus genomic separation for plants and mammals (top) with an $s^{-1}$ reference (dashed). The lower subpanel shows the log–log derivative ($d\log P/\, d\log s$) with a guide at ~40 kb (magenta). Human and mouse ESCs display a shoulder at ~$10^5$–$10^6$ bp and a corresponding rise in the derivative, consistent with barrier-stalled loop extrusion; plant profiles lack a pronounced shoulder and follow a smoother $\approx s^{-1}$ decay. B. Distributions of genomic distances between adjacent insulation-valley (IV) centers (top) and chromatin-loop sizes (bottom). c. Summary statistics for IV spacing and loop sizes across species. All datasets were down-sampled to comparable read depth before IV and loop detection; therefore, absolute counts differ slightly from those obtained when only *S.lycopersicum* and mESC were depth-normalized.

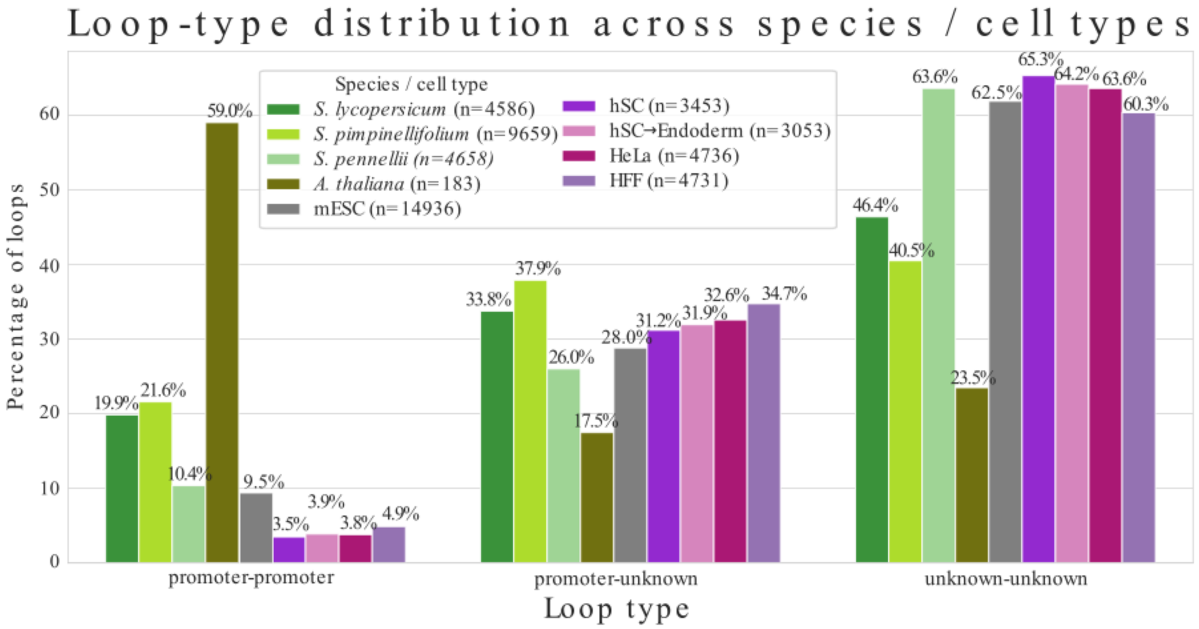


Supplementary Figure 16: **Distributions of chromatin loop types across plant species and mammalian cells.** The total number of detected loops is indicated in parentheses.

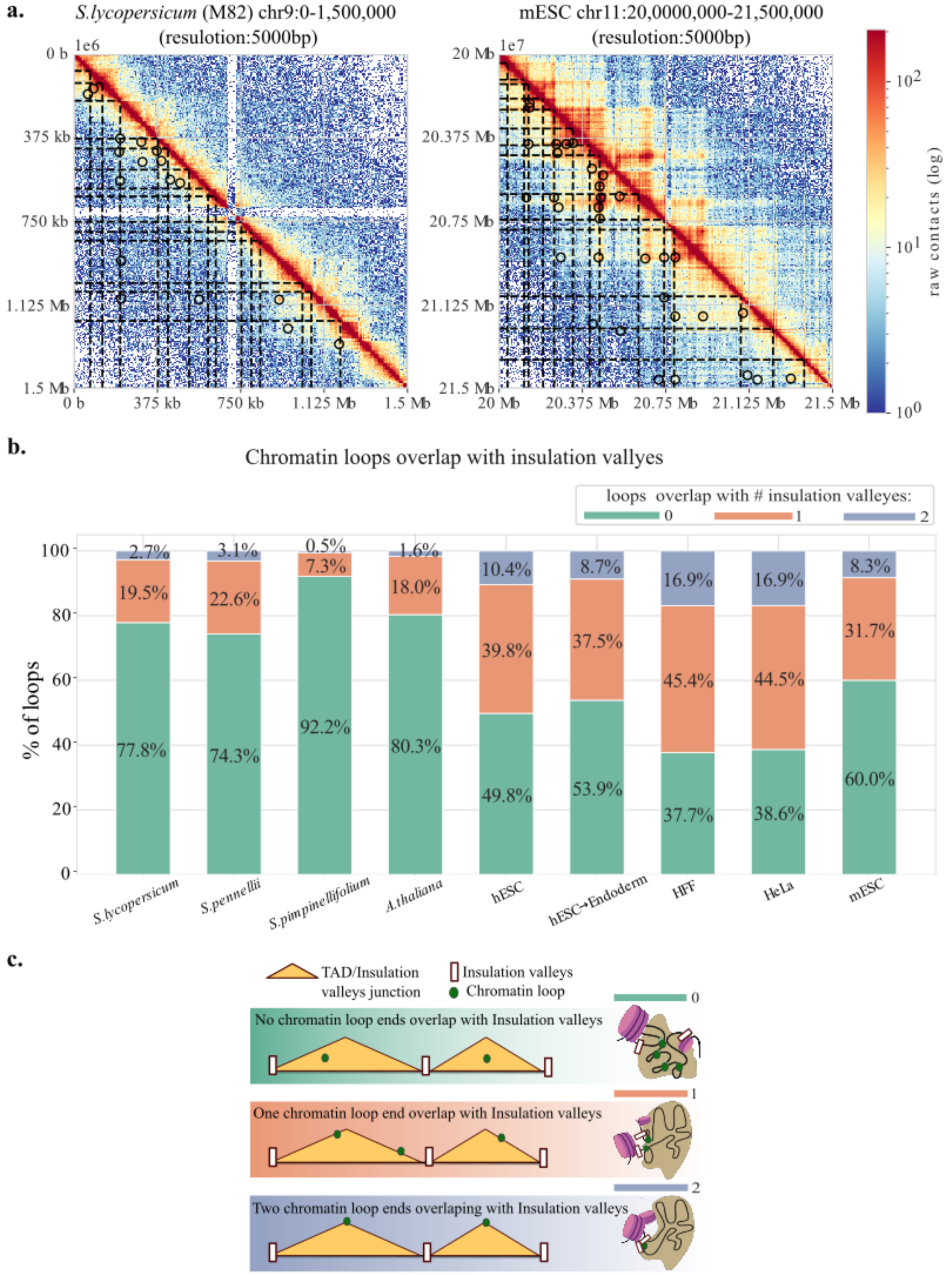


Supplementary Figure 17: **Chromatin loop overlap with insulation valleys across plant species and mammalian cells.** a. Heatmaps of *S.lycopersicum* (left) and mESC cells (right), with insulation-valley centers marked by black dashed lines and chromatin loops indicated by black circles. b. Proportion of chromatin loops that overlap insulation-valley centers. c. Schematic models illustrating the different overlap scenarios.

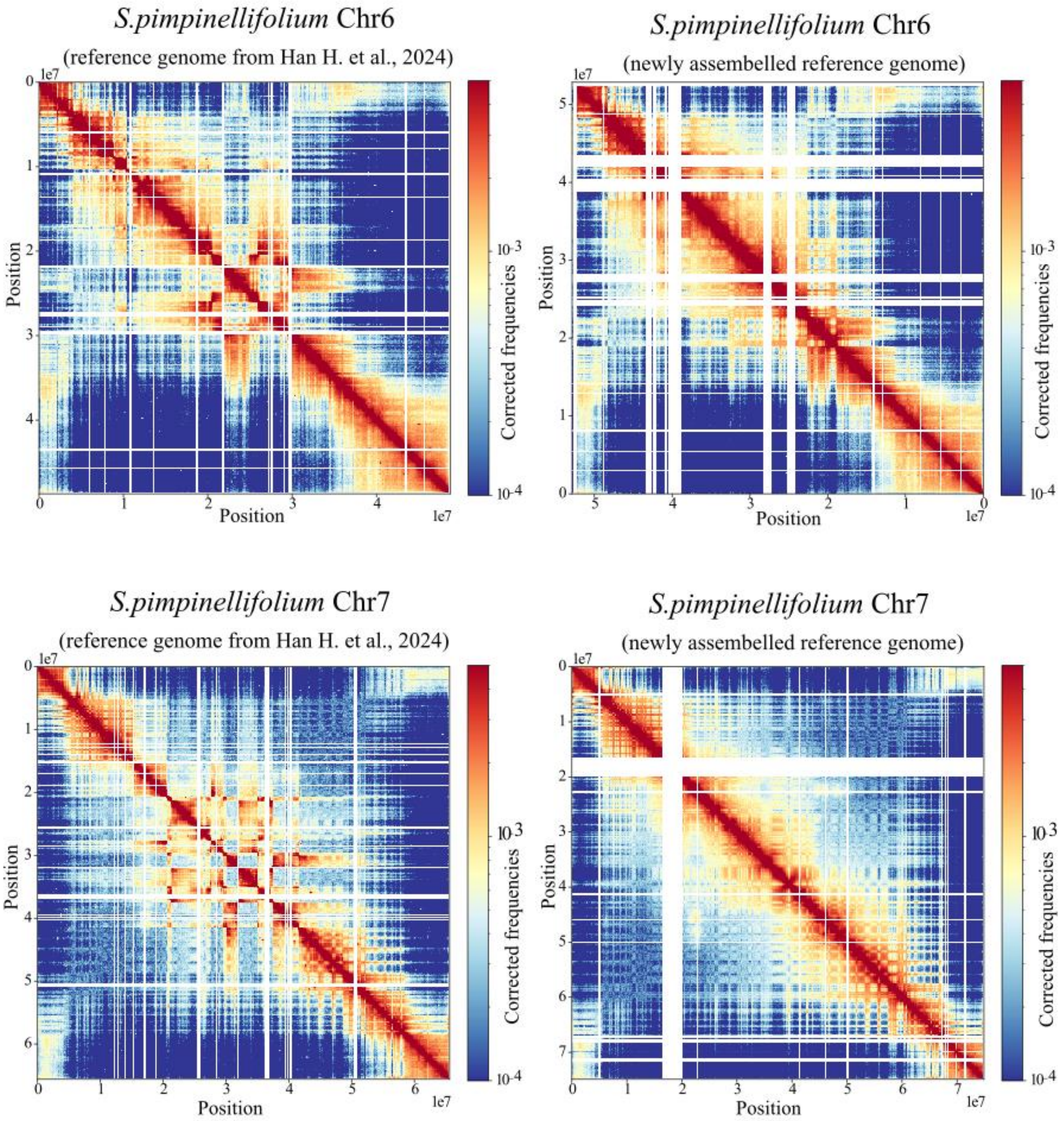


Supplementary Figure 18: **Micro-C contact maps for *S.pimpinellifolium* generated with two LA1589 references: the published assembly[68] (left) and a new assembly built from the specific LA1589 seed stock used in this study (right).** Heatmaps are shown at 100-kb resolution with an identical color scale. Divergent interaction patterns highlight lineage-specific chromosomal rearrangements that become apparent only when the Micro-C data are aligned to a reference derived from the same seed source.

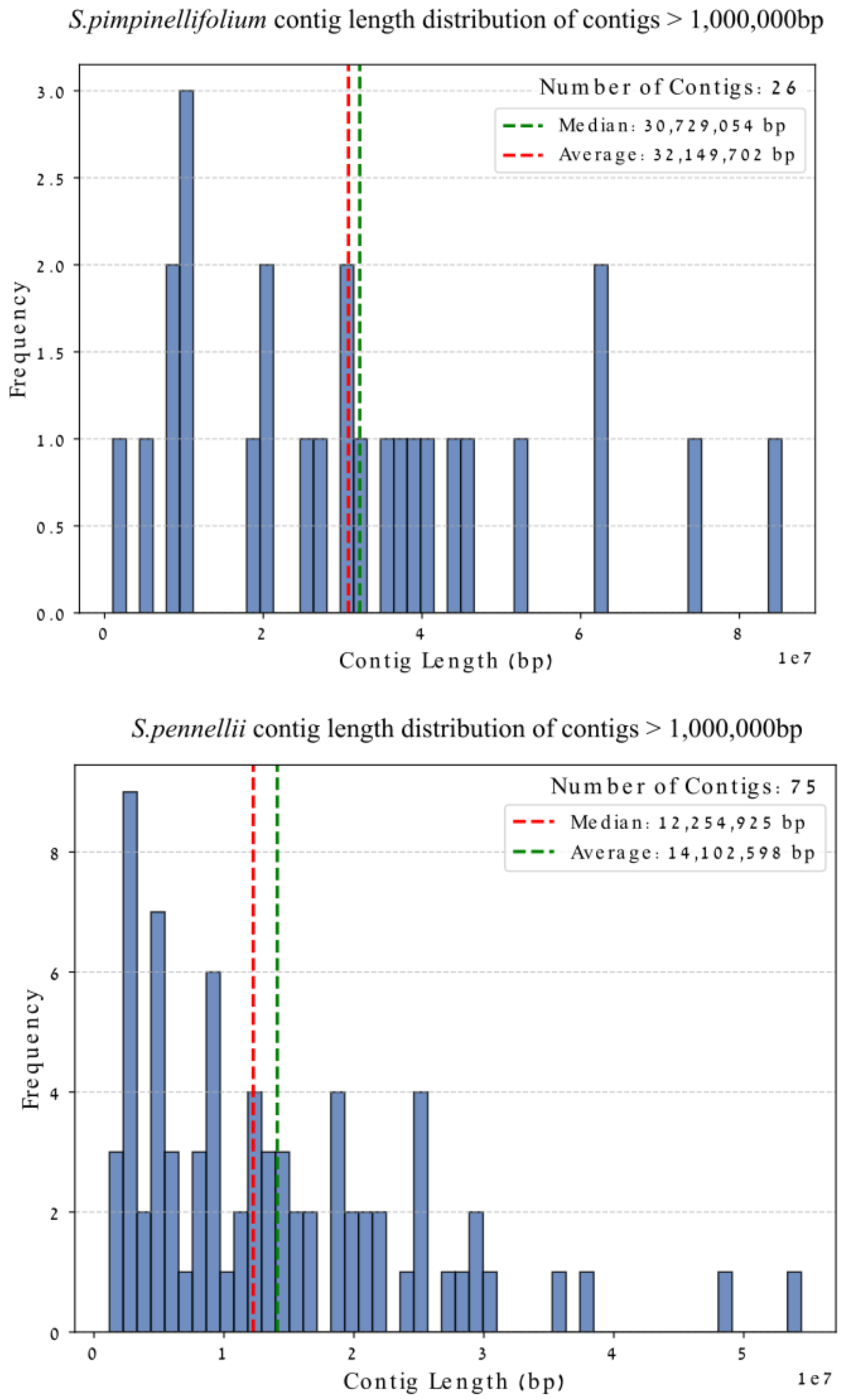


Supplementary Figure 19: C**ontig length distributions (> 1 Mb) for *S.pimpinellifolium* and *S.pennellii.***

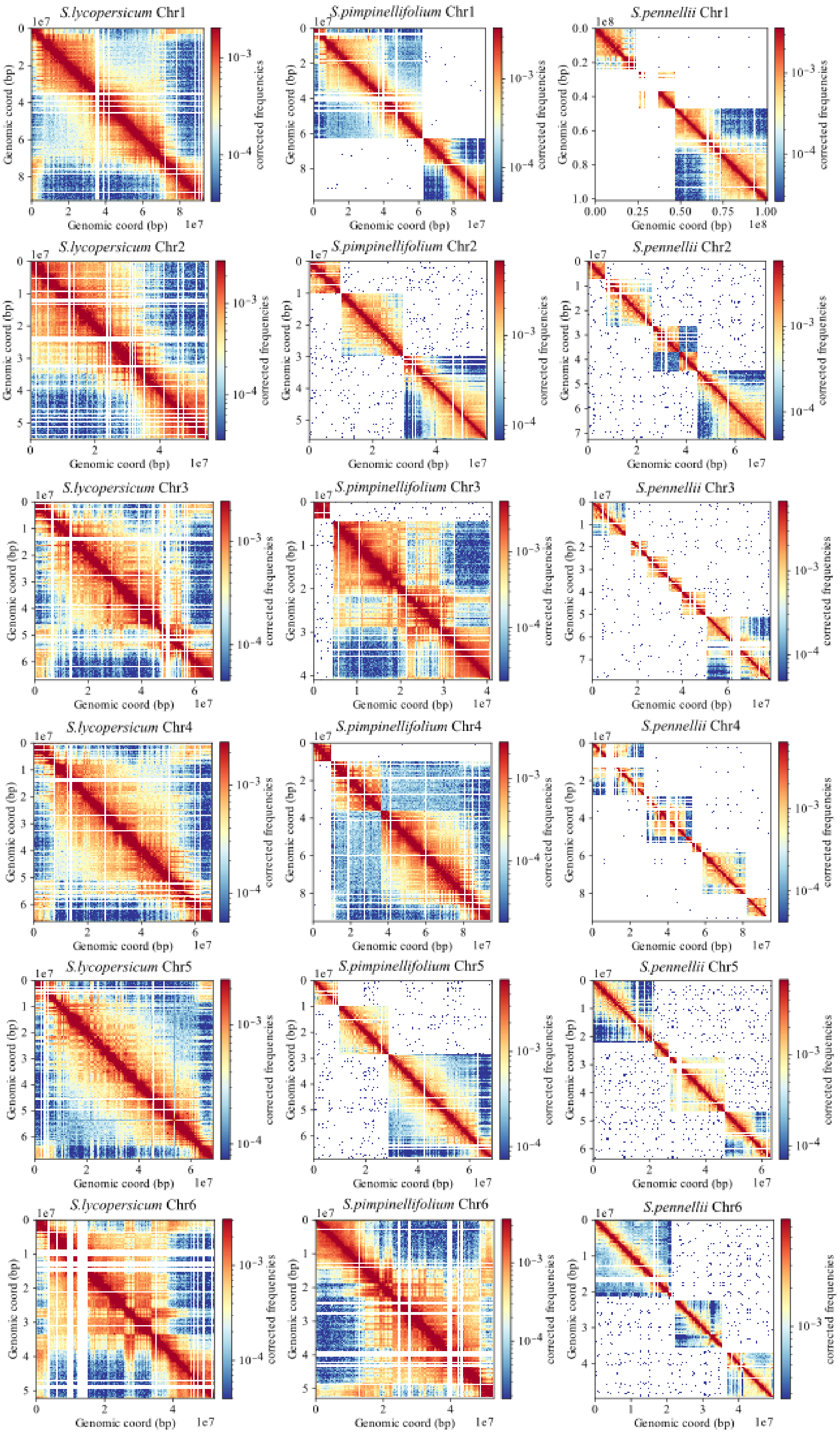

*S.lycopersicum* Chr1
*S.pimpinellifolium* Chr1
*S.pennellii* Chr1
*S.lycopersicum* Chr2
*S.pimpinellifolium* Chr2
*S.pennellii* Chr2
*S.lycopersicum* Chr3
*S.pimpinellifolium* Chr3
*S.pennellii* Chr3
*S.lycopersicum* Chr4
*S.pimpinellifolium* Chr4
*S.pennellii* Chr4
*S.lycopersicum* Chr5
*S.pimpinellifolium* Chr5
*S.pennellii* Chr5
*S.lycopersicum* Chr6
*S.pimpinellifolium* Chr6
*S.pennellii* Chr6
Genomic coord (bp)
corrected frequencies

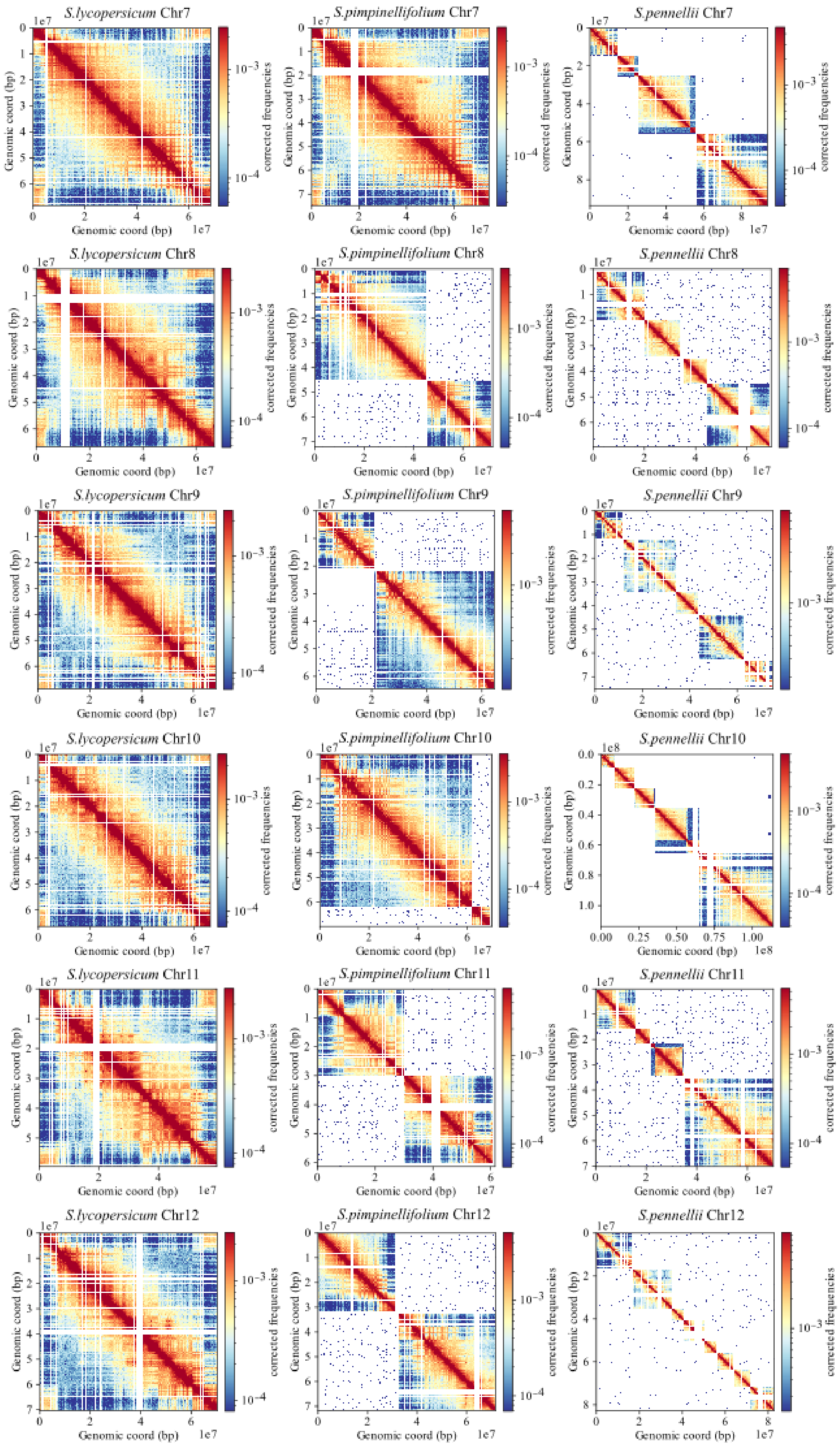

S.lycopersicum Chr7
S.pimpinellifolium Chr7
S.pennellii Chr7
S.lycopersicum Chr8
S.pimpinellifolium Chr8
S.pennellii Chr8
S.lycopersicum Chr9
S.pimpinellifolium Chr9
S.pennellii Chr9
S.lycopersicum Chr10
S.pimpinellifolium Chr10
S.pennellii Chr10
S.lycopersicum Chr11
S.pimpinellifolium Chr11
S.pennellii Chr11
S.lycopersicum Chr12
S.pimpinellifolium Chr12
S.pennellii Chr12
Genomic coord (bp)
corrected frequencies
1e7
1e8

### Whole Chromosome Similarity Metrics

| Chromosome | Comparison | Pearson | Spearman | MSE | SSIM |
|---|---|---|---|---|---|
| chr1 | M82 vs. *S.pimpinellifolium* | 0.975 | 0.688 | 0.0 | 0.995 |
| chr1 | M82 vs. *S.pennellii* | 0.898 | 0.49 | 0.0 | 0.994 |
| chr2 | M82 vs. *S.pimpinellifolium* | 0.975 | 0.642 | 0.0 | 0.987 |
| chr2 | M82 vs. *S.pennellii* | 0.836 | 0.636 | 0.0 | 0.99 |
| chr3 | M82 vs. *S.pimpinellifolium* | 0.968 | 0.73 | 0.0 | 0.988 |
| chr3 | M82 vs. *S.pennellii* | 0.651 | 0.518 | 0.001 | 0.981 |
| chr4 | M82 vs. *S.pimpinellifolium* | 0.966 | 0.526 | 0.0 | 0.993 |
| chr4 | M82 vs. *S.pennellii* | 0.677 | 0.597 | 0.001 | 0.986 |
| chr5 | M82 vs. *S.pimpinellifolium* | 0.973 | 0.742 | 0.0 | 0.985 |
| chr5 | M82 vs. *S.pennellii* | 0.815 | 0.593 | 0.0 | 0.99 |
| chr6 | M82 vs. *S.pimpinellifolium* | 0.904 | 0.454 | 0.0 | 0.992 |
| chr6 | M82 vs. *S.pennellii* | 0.799 | 0.587 | 0.0 | 0.993 |
| chr7 | M82 vs. *S.pimpinellifolium* | 0.963 | 0.773 | 0.0 | 0.994 |
| chr7 | M82 vs. *S.pennellii* | 0.887 | 0.648 | 0.0 | 0.991 |
| chr8 | M82 vs. *S.pimpinellifolium* | 0.966 | 0.655 | 0.0 | 0.995 |
| chr8 | M82 vs. *S.pennellii* | 0.713 | 0.588 | 0.001 | 0.989 |
| chr9 | M82 vs. *S.pimpinellifolium* | 0.962 | 0.661 | 0.0 | 0.994 |
| chr9 | M82 vs. *S.pennellii* | 0.702 | 0.625 | 0.001 | 0.986 |
| chr10 | M82 vs. *S.pimpinellifolium* | 0.972 | 0.777 | 0.0 | 0.994 |
| chr10 | M82 vs. *S.pennellii* | 0.837 | 0.692 | 0.0 | 0.987 |
| chr11 | M82 vs. *S.pimpinellifolium* | 0.909 | 0.549 | 0.0 | 0.995 |
| chr11 | M82 vs. *S.pennellii* | 0.871 | 0.638 | 0.0 | 0.991 |
| chr12 | M82 vs. *S.pimpinellifolium* | 0.965 | 0.63 | 0.0 | 0.993 |
| chr12 | M82 vs. *S.pennellii* | 0.747 | 0.596 | 0.001 | 0.986 |

Supplementary Figure 20: **Comparison of Micro-C heatmaps across all chromosomes in *S.lycopersicum* vs. *S.pimpinellifolium* and *S.pennellii.*** Each panel corresponds to one chromosome. The accompanying table summarizes Pearson's and Spearman's correlation coefficients, Mean Squered Error (MSE) and Structural Similarity Index (SSIM) for each chromosome at the whole-chromosome level.

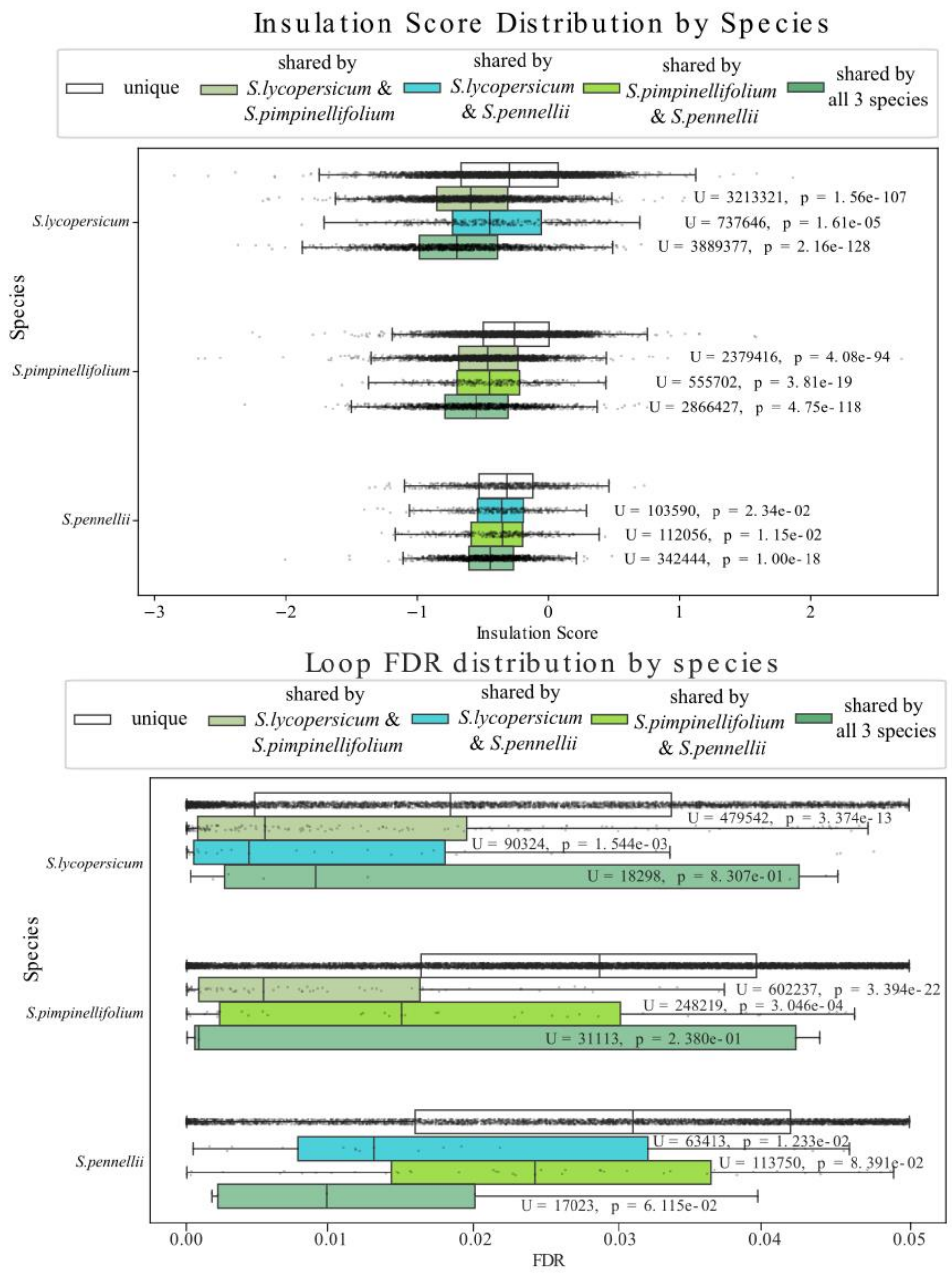


Supplementary Figure 21**: Distribution of insulation scores and loop FDRs across the three *Solanum* species.** Each point is an individual insulation valley (IV) score (upper panel) or loop FDR (lower panel). Box-plots summarise five classes: unique, shared by *S.lycopersicum* & *S.pimpinellifolium*, shared by *S.lycopersicum* & *S.pennellii*, shared by *S.pimpinellifolium* & *S.pennellii*, and shared by all three species. For every species the distribution of each shared class was compared with that species' unique class using a two-sided Mann-Whitney U test; the U statistic and P-value are printed beside the corresponding box.

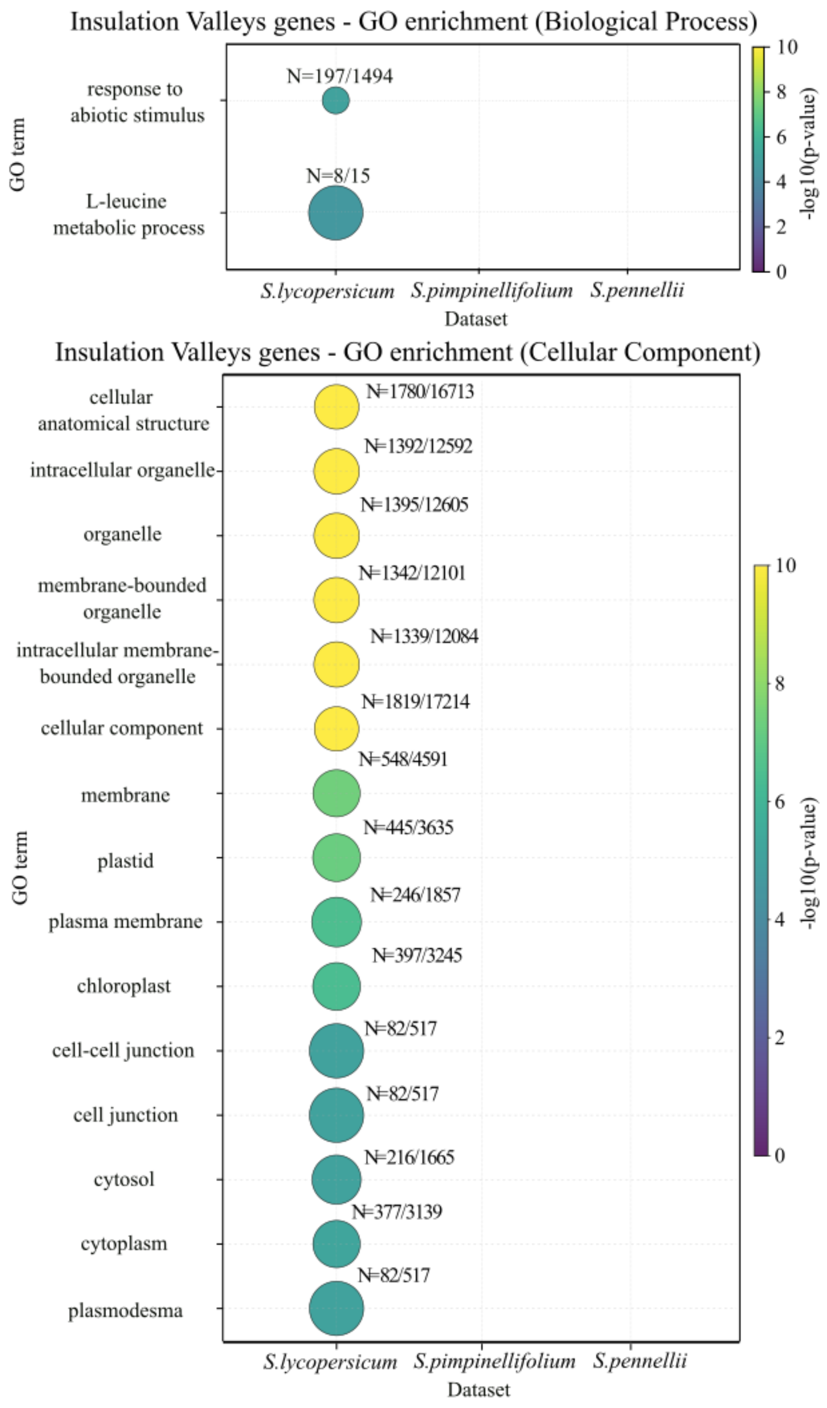


Supplementary Figure 22**: GO enrichment of Insulation Valleys genes in *S.lycopersicum* compared with *S.pimpinellifolium* and *S.pennellii*.** Dot size denotes the fraction of IV-associated genes within each GO term (N = IV genes in term/total genes assigned to that term). Dot colour reflects the -log10 p-value of the enrichment test; lighter colors indicate stronger significance

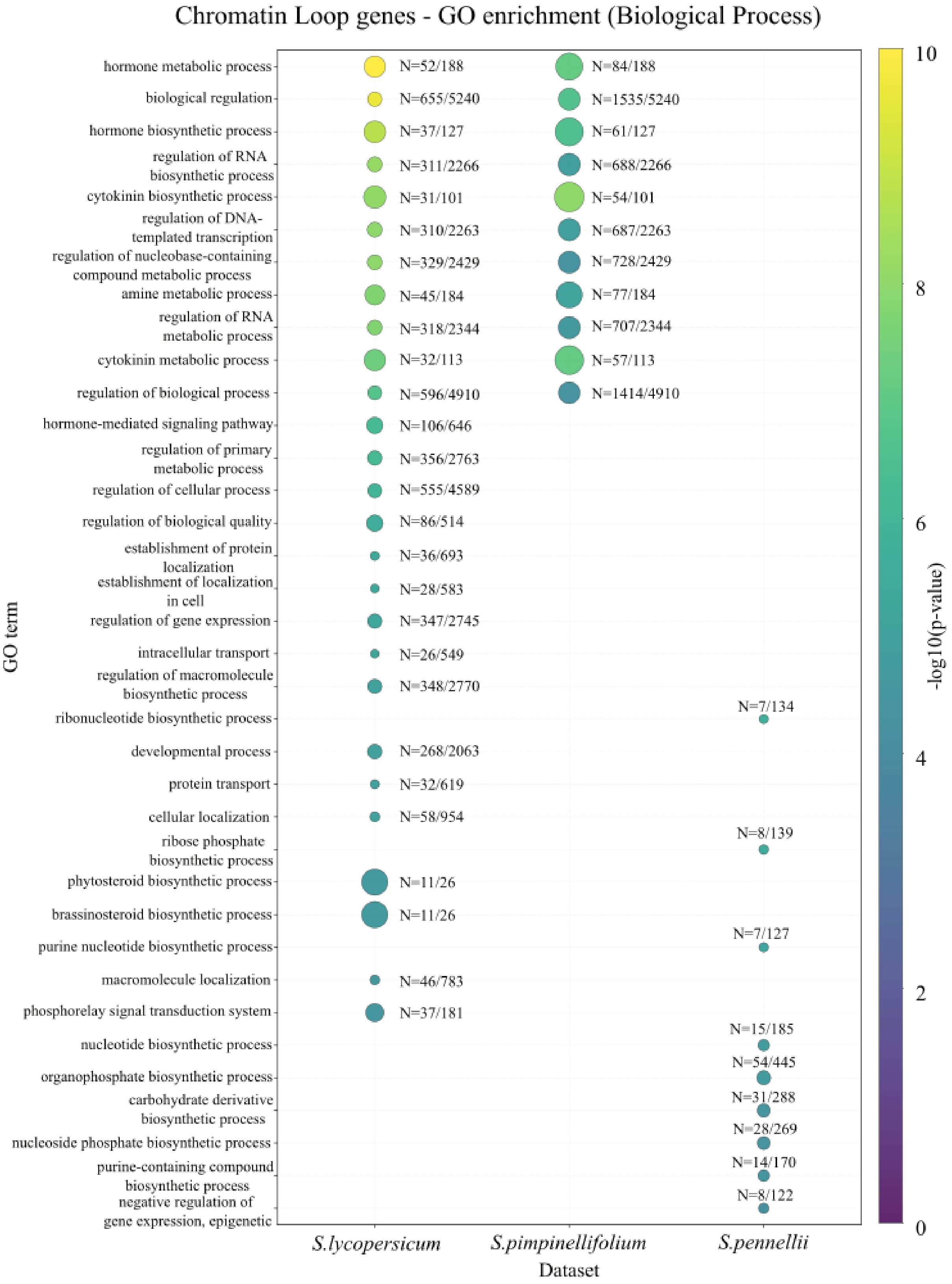
Chromatin Loop genes - GO enrichment (Biological Process)
GO term
hormone metabolic process
biological regulation
hormone biosynthetic process
regulation of RNA biosynthetic process
cytokinin biosynthetic process
regulation of DNA-templated transcription
regulation of nucleobase-containing compound metabolic process
amine metabolic process
regulation of RNA metabolic process
cytokinin metabolic process
regulation of biological process
hormone-mediated signaling pathway
regulation of primary metabolic process
regulation of cellular process
regulation of biological quality
establishment of protein localization
establishment of localization in cell
regulation of gene expression
intracellular transport
regulation of macromolecule biosynthetic process
ribonucleotide biosynthetic process
developmental process
protein transport
cellular localization
ribose phosphate biosynthetic process
phytosteroid biosynthetic process
brassinosteroid biosynthetic process
purine nucleotide biosynthetic process
macromolecule localization
phosphorelay signal transduction system
nucleotide biosynthetic process
organophosphate biosynthetic process
carbohydrate derivative biosynthetic process
nucleoside phosphate biosynthetic process
purine-containing compound biosynthetic process
negative regulation of gene expression, epigenetic
N=52/188
N=655/5240
N=37/127
N=311/2266
N=31/101
N=310/2263
N=329/2429
N=45/184
N=318/2344
N=32/113
N=596/4910
N=106/646
N=356/2763
N=555/4589
N=86/514
N=36/693
N=28/583
N=347/2745
N=26/549
N=348/2770
N=268/2063
N=32/619
N=58/954
N=11/26
N=11/26
N=46/783
N=37/181
N=84/188
N=1535/5240
N=61/127
N=688/2266
N=54/101
N=687/2263
N=728/2429
N=77/184
N=707/2344
N=57/113
N=1414/4910
N=7/134
N=8/139
N=7/127
N=15/185
N=54/445
N=31/288
N=28/269
N=14/170
N=8/122
*S.lycopersicum*
*S.pimpinellifolium*
*S.pennellii*
Dataset
-log10(p-value)
0
2
4
6
8
10

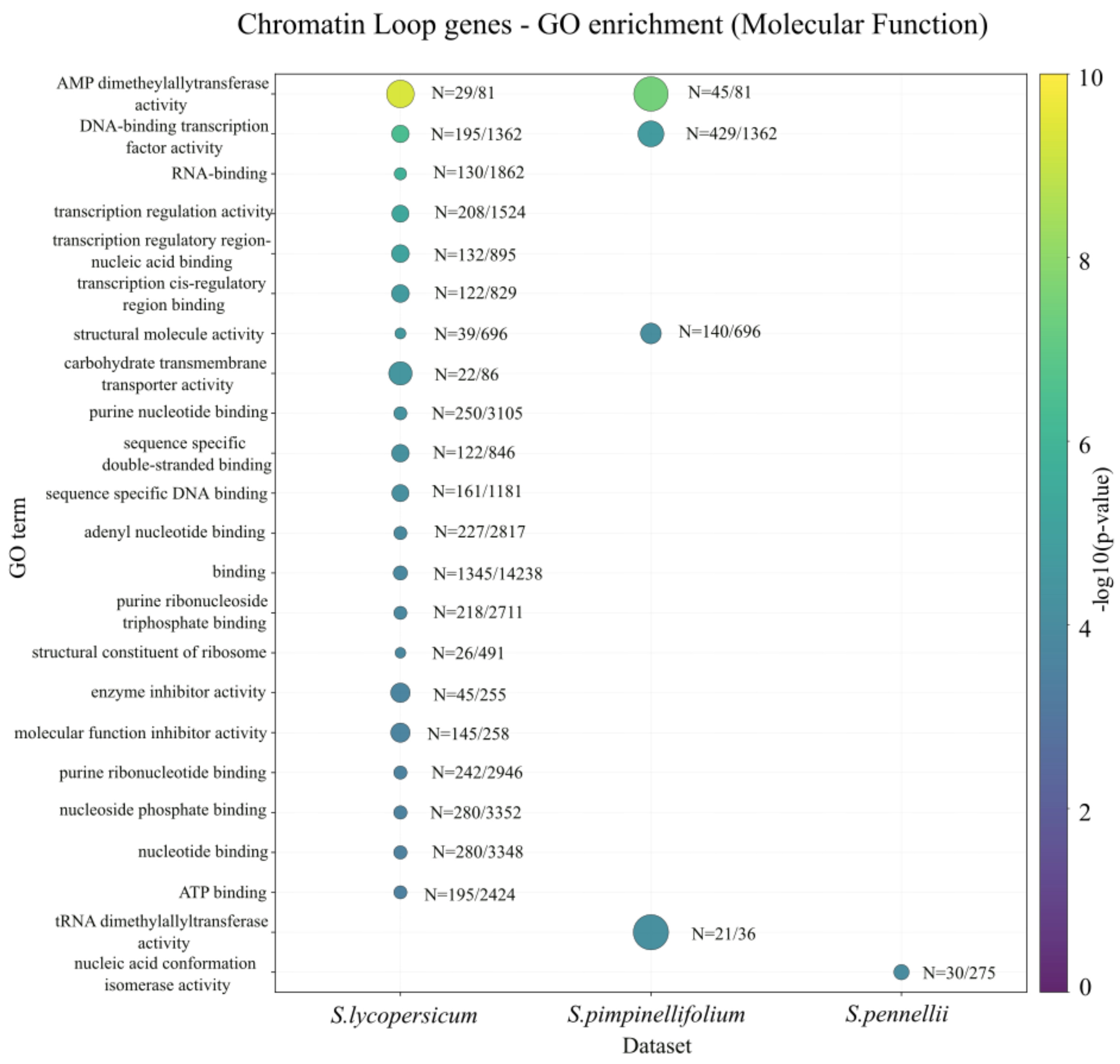
Chromatin Loop genes - GO enrichment (Molecular Function)
AMP dimetheylallyltransferase activity
DNA-binding transcription factor activity
RNA-binding
transcription regulation activity
transcription regulatory region-nucleic acid binding
transcription cis-regulatory region binding
structural molecule activity
carbohydrate transmembrane transporter activity
purine nucleotide binding
sequence specific double-stranded binding
sequence specific DNA binding
adenyl nucleotide binding
binding
purine ribonucleoside triphosphate binding
structural constituent of ribosome
enzyme inhibitor activity
molecular function inhibitor activity
purine ribonucleotide binding
nucleoside phosphate binding
nucleotide binding
ATP binding
tRNA dimethylallyltransferase activity
nucleic acid conformation isomerase activity
GO term
N=29/81
N=45/81
N=195/1362
N=429/1362
N=130/1862
N=208/1524
N=132/895
N=122/829
N=39/696
N=140/696
N=22/86
N=250/3105
N=122/846
N=161/1181
N=227/2817
N=1345/14238
N=218/2711
N=26/491
N=45/255
N=145/258
N=242/2946
N=280/3352
N=280/3348
N=195/2424
N=21/36
N=30/275
S.lycopersicum
S.pimpinellifolium
S.pennellii
Dataset
-log10(p-value)
0
2
4
6
8
10

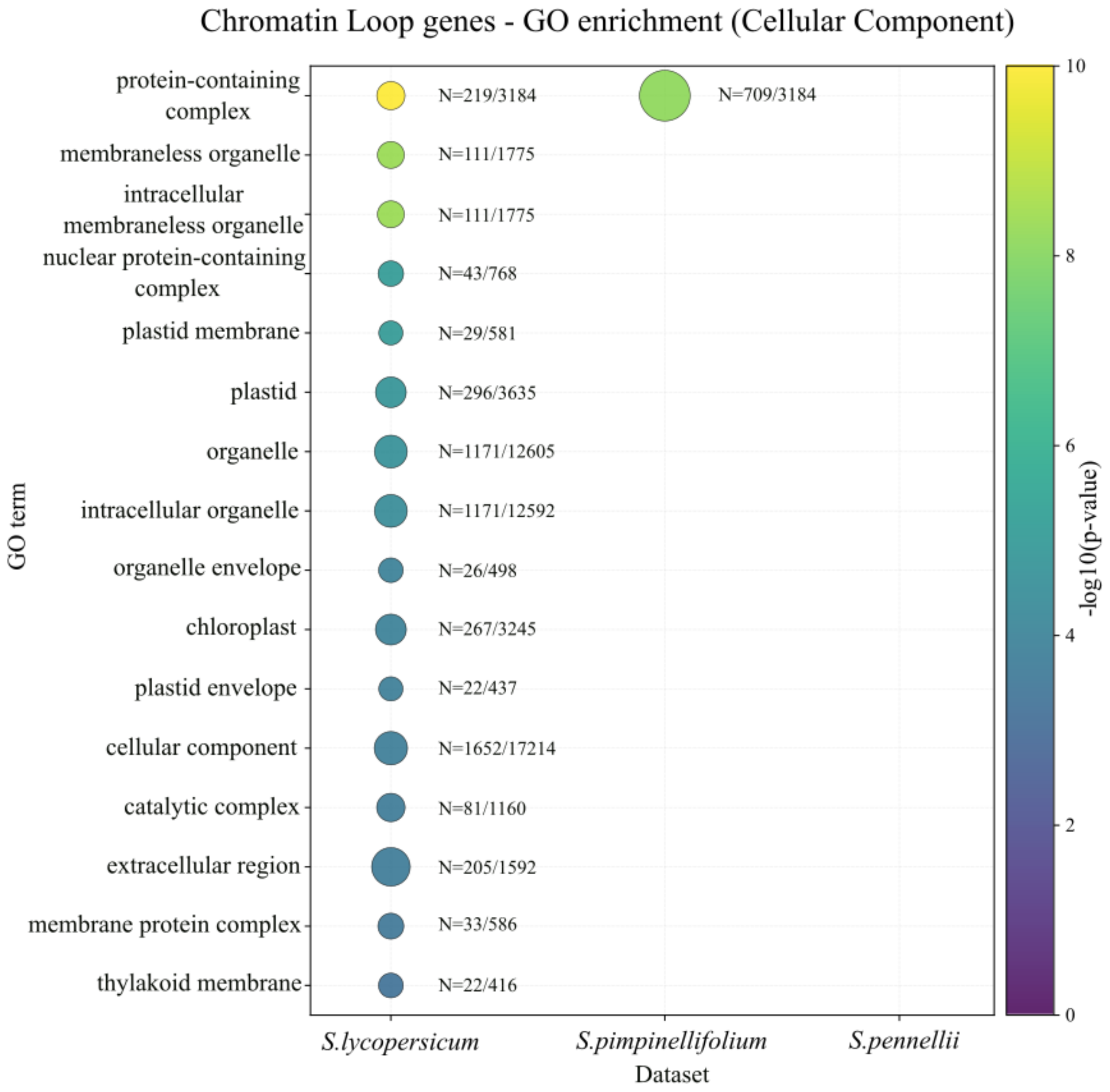


Supplementary Figure 23**: GO enrichment of chromatin loop genes in *S.lycopersicum* compared with *S.pimpinellifolium* and *S.pennellii*.** Dot size denotes the fraction of IV-associated genes within each GO term (N = IV genes in term/total genes assigned to that term). Dot color reflects the -log10 p-value of the enrichment test; lighter colors indicate stronger significance.

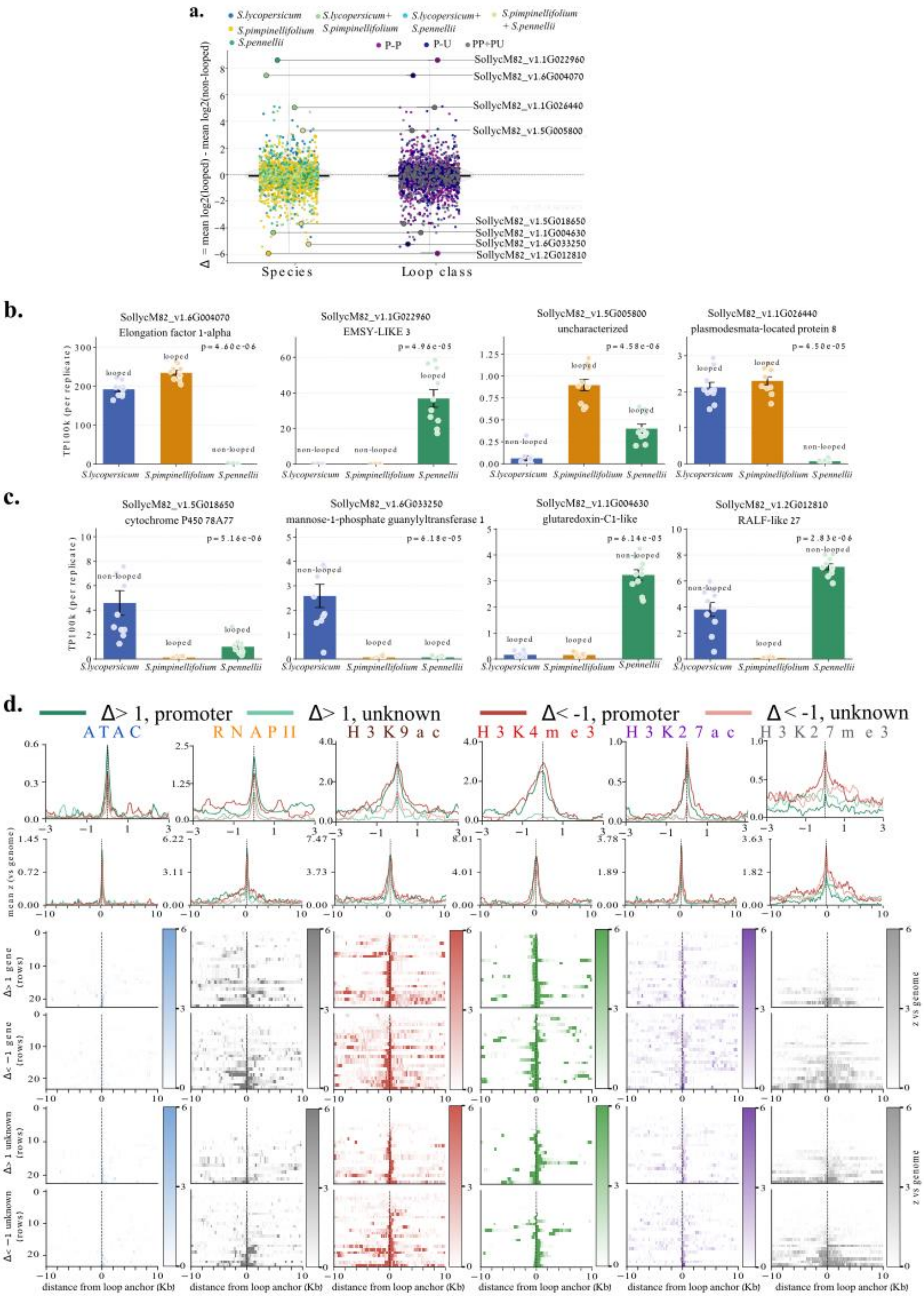

a.
S.lycopersicum
S.lycopersicum+ S.pimpinellifolium
S.lycopersicum+ S.pennellii
S.pimpinellifolium + S.pennellii
S.pimpinellifolium
S.pennellii
P-P
P-U
PP+PU
Δ = mean log2(looped) - mean log2(non-looped)
SollycM82_v1.1G022960
SollycM82_v1.6G004070
SollycM82_v1.1G026440
SollycM82_v1.5G005800
SollycM82_v1.5G018650
SollycM82_v1.1G004630
SollycM82_v1.6G033250
SollycM82_v1.2G012810
Species
Loop class
b.
SollycM82_v1.6G004070
Elongation factor 1-alpha
p = 4.60e-06
SollycM82_v1.1G022960
EMSY-LIKE 3
p = 4.96e-05
SollycM82_v1.5G005800
uncharacterized
p = 4.58e-06
SollycM82_v1.1G026440
plasmodesmata-located protein 8
p = 4.50e-05
TP100k (per replicate)
looped
non-looped
S.lycopersicum
S.pimpinellifolium
S.pennellii
c.
SollycM82_v1.5G018650
cytochrome P450 78A77
p = 5.16e-06
SollycM82_v1.6G033250
mannose-1-phosphate guanylyltransferase 1
p = 6.18e-05
SollycM82_v1.1G004630
glutaredoxin-C1-like
p = 6.14e-05
SollycM82_v1.2G012810
RALF-like 27
p = 2.83e-06
d.
Δ> 1, promoter
Δ> 1, unknown
Δ< -1, promoter
Δ< -1, unknown
ATAC
RNAPII
H3K9ac
H3K4me3
H3K27ac
H3K27me3
mean z (vs genome)
Δ> 1 gene (rows)
Δ< −1 gene (rows)
Δ> 1 unknown (rows)
Δ< −1 unknown (rows)
z vs genome
distance from loop anchor (Kb)

Supplementary Figure 24: **Cross-species loop-expression contrast (Δ), example genes, and anchor-centered chromatin profiles.** a. Genome-wide Δ distribution- For each gene expressed in all three species, the cross-species contrast was computed as $\Delta = \text{mean}[\log_2(\text{TP100k})]_{\text{looped}} - \text{mean}[\log_2(\text{TP100k})]_{\text{non-looped}}$. Left: Δ values by species in which the promoter is looped; right: the same Δ values grouped by loop class (PP, PU, or genes participating in both). Points are individual genes; outliers are labeled. The distribution is continuous, spanning strongly positive to strongly negative values. b. Representative genes with $\Delta > 1$ (enhanced when looped) - Per species, bars summarize TP100k for looped versus non-looped states; dots are biological replicates. Error bars denote dispersion. Two-sided Mann-Whitney U p-values (looped vs non-looped) are shown above panels. c. Representative genes with $\Delta < -1$ (repressed/poised when looped) - Same display as in b, illustrating genes whose expression is lower in looped species than in non-looped species. d. Anchor-centered chromatin at P–U loops stratified by Δ and anchor type**-** For ATAC, RNAPII, H3K9ac, H3K4me3, H3K27ac, and H3K27me3, the top traces show the mean signal (z-score vs a genome background) and the heatmaps below show individual loci, all aligned at the loop anchor (dashed line) within ±3 kb (top panel) and ±10 kb (bottom panel). Columns are marks; groups indicate $\Delta > 1$ or $\Delta < -1$ and whether the profiled anchor is the promoter or the unknown partner. Activating marks (ATAC, RNAPII, H3K9ac, H3K27ac) peak at promoters for both Δ groups and are weaker at unknown anchors, whereas H3K27me3 is specifically elevated for $\Delta < -1$ at both the promoter and the distal (unknown) anchor. Color bars denote z-score scale.

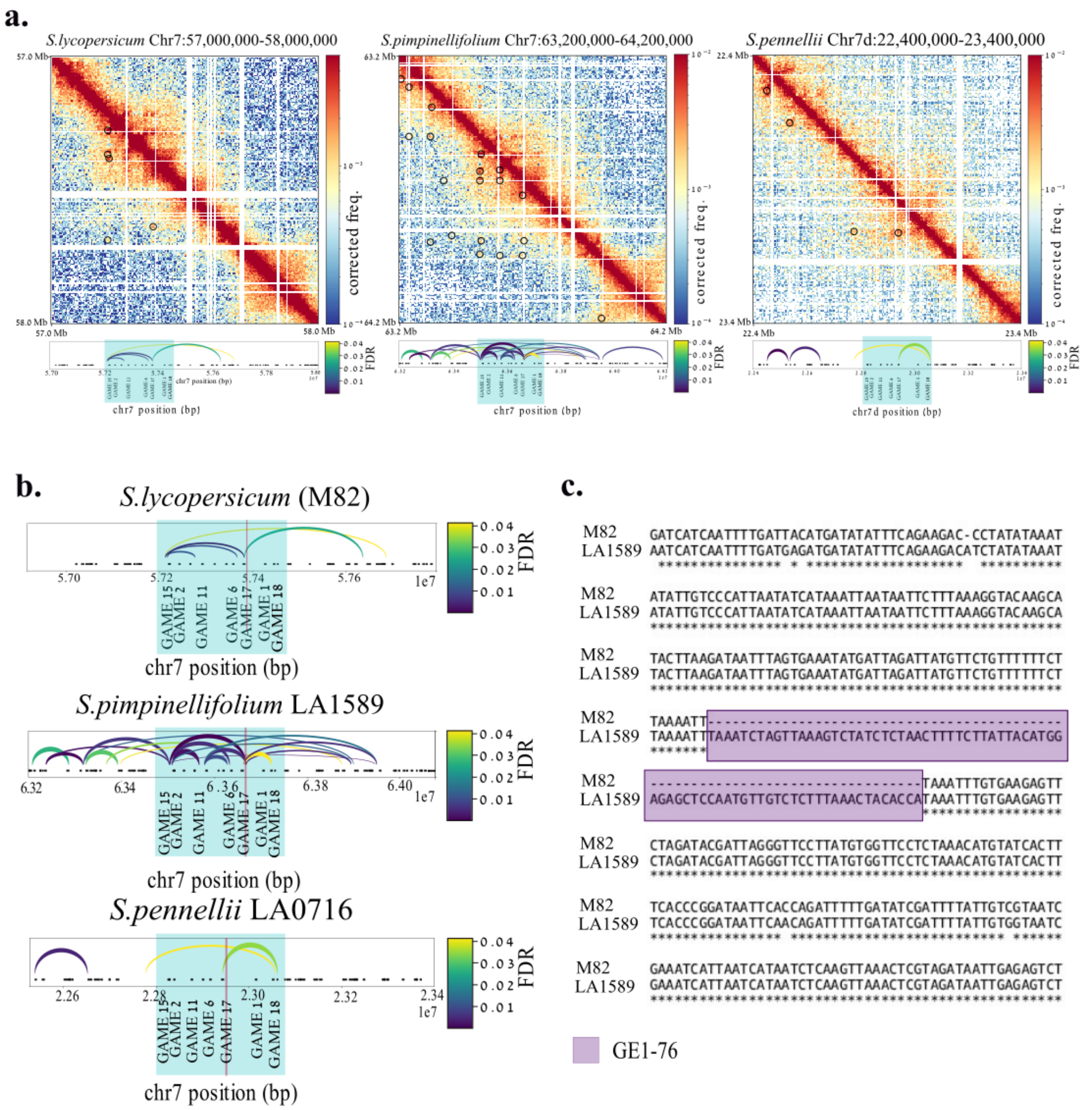


Supplementary Figure 25: **Chromatin looping and sequence variation at the GAME Enhancer 1 (GE1) locus.** a. Micro-C contact maps (5 kb resolution) for the GAME1 region on chromosome 7 in *S.lycopersicum* (M82), *S.pimpinellifolium* (LA1589) and *S.pennellii* (LA0716). black circles mark loops called by Mustache (FDR < 0.05). Below each map, loop-arc diagrams plot all significant interactions (color = FDR; legend right), with the GAME gene cluster. b. Zoomed-in loop-arc plots for each species, centered on the GE1 enhancer region (shaded box). Loop anchors are labeled by GAME gene ID; the red line repressent GE1-the GAME identified enhancer. C. Pairwise nucleotide alignment of the GE1 interval between *S.lycopersicum* (M82) and *S.pimpinellifolium* (LA1589). Asterisks denote identical bases; the 76-bp polymorphic element (“GE1-76”) present in LA1589 is highlighted in purple.

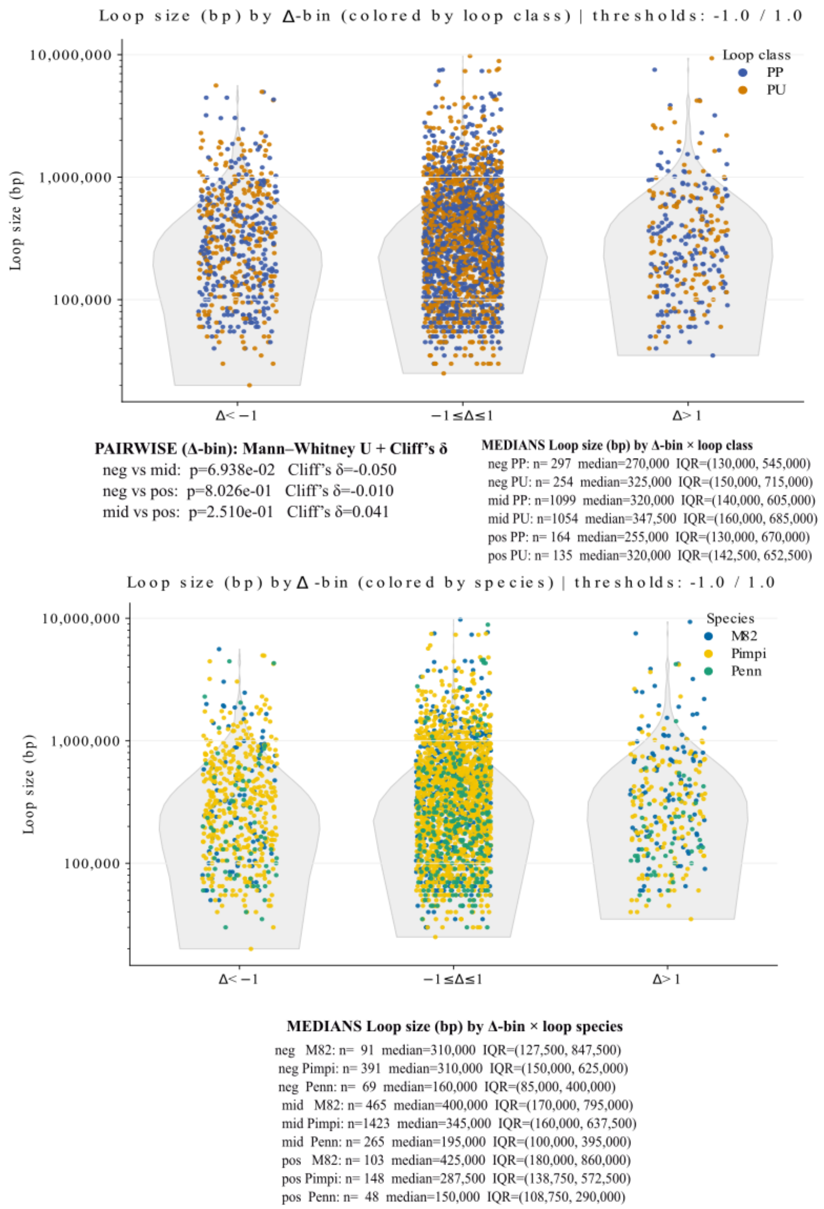


Supplementary Figure 26: **Loop-size does not predict enhancement or repression, and patterns are independent of loop class and species.**

Loops were assigned a cross-species expression contrast $\Delta = \text{mean}[\log_2(\text{TP100k})]_{\text{looped}} - \text{mean}[\log_2(\text{TP100k})]_{\text{non-looped}}$. Loops were restricted to cis PP or PU and sized as the genomic distance between anchor midpoints ($\log_{10}$ y-axis). Δ bins: $\Delta < -1$ (repressed), $-1 \leq \Delta \leq 1$ (neutral), $\Delta > 1$ (enhanced).

**Top:** Violin plots of loop size colored by loop class (PP blue, PU orange) with jittered points. Across Δ bins, loop-size distributions are similar (medians ≈270-347 kb; see inset). Pairwise Mann-Whitney tests between bins are non-significant after Bonferroni correction and effect sizes are negligible (|Cliff's δ|≤0.05). Composition of PP vs PU does not vary with Δ (2×3 $\chi^2$ p=0.28; Cramér's V=0.03).

**Bottom:** The same violins colored by species (M82, *S. pimpinellifolium*, *S. pennellii*). Median sizes differ only modestly across species within each Δ bin (see inset), and no species shows a systematic shift with Δ.
